# Cryo-electron tomography of Nipah virus structural protein complexes in virus-like particles

**DOI:** 10.64898/2026.07.16.738223

**Authors:** Viraj V. Upadhye, Jean F. Lee, Nihan Ercanli, Clifton Ricana, Amy C. Hinsley, Martin Obr, Ludovic Autin, Florian K. M. Schur, Hector C. Aguilar, Robert A. Dick

## Abstract

Nipah virus (NiV) is a BSL-4 zoonotic paramyxovirus with ∼75% human mortality. The matrix protein (M) of NiV and other paramyxoviruses binds the inner leaflet of the cellular plasma membrane, orchestrating virion assembly by bringing together transmembrane glycoproteins (F/G) and ribonucleoprotein complexes (N). However, the interactions of these full-length proteins within membrane complexes remain elusive. Using cryo-electron tomography and subtomogram averaging of virus like particles (VLPs), we interrogated the protein:protein interactions of the main NiV structural proteins M/N/F/G. The M lattice structure determined to 7Å revealed a novel M-dimer arrangement that yielded two distinct repeating holes. Notably, F-trimers were arranged above only one of the two holes, dependent on F’s cytoplasmic tail. G was enriched in regions of higher M-VLP curvature, while N dramatically increased M-VLP pleomorphism. This work provides novel insights into paramyxoviral protein complexes, structures, and morphology.

## Introduction

Paramyxoviruses are pervasive human pathogens that include measles virus (MeV), mumps virus (MuV), human parainfluenza viruses (hPIVs), and the deadly zoonotic henipaviruses Nipah (NiV) and Hendra (HeV) viruses ^1–3^. Henipaviruses can spread from bats to humans, as well as from human to human, and such zoonotic spread from animals to humans is projected to increase in the future ^4–8^. No approved human vaccines or therapeutics exist for henipaviruses, highlighting a need for medical intervention that could be quickly implemented if a pandemic arose ^9^. Despite their relevance to human health, due to their high-containment biosafety level 4 (BSL-4) requirements and select-agent status, structural studies of authentic henipavirus particles have not been feasible.

We and others have shown that expression of some paramyxoviral structural proteins in mammalian cells is sufficient for assembly and budding of virus-like particles (VLPs) ^10–20^. These VLPs assemble autonomously, in complex with the cellular membrane, and are released into the medium of transfected cells. VLP systems provide an excellent platform to study the full-length near-native assemblies of highly pathogenic paramyxoviruses, such as NiV, in the absence of high containment. While cryo-electron microscopy (cryoEM) has been used to study the structures of some infectious paramyxoviruses [MeV, and Newcastle Disease virus (NDV)], pneumoviruses, [respiratory syncytial virus (RSV)], and orthomyxoviruses, (influenza viruses), this study is the first to leverage VLPs for sub-nanometer structure determination of a paramyxovirus by cryoEM ^12,21–23^. NiV encodes four major structural proteins: nucleocapsid (N), matrix (M), fusion (F), and receptor binding (G), as well as two nonstructural proteins, P and L. M is the master assembly protein found in all paramyxoviruses, and it recruits both viral and cellular proteins used for viral assembly and egress ^24,25^. Expression of M alone in mammalian cells is sufficient for assembly and budding of VLPs, which are roughly spherical in morphology when imaged by negative stain transmission electron-microscopy (nsTEM) ^11,13,15,16,18,26–28^. F and G are transmembrane glycoproteins found on the surface of viral particles and are required for host cell recognition and viral entry. F and G are promising targets for use in vaccines and therapeutics due to their inherent immunogenicity, but their structures and organization relative to the viral envelope and other viral proteins are not well understood ^29–31^. We and others have shown that expression of some paramyxovirus F proteins can result in VLP formation ^19,32^. N is required for the incorporation of the viral genome into progeny virions and also acts as a transcriptional regulator ^14,33–35^.

Purified soluble versions of the four main NiV structural proteins (herein defined as N/M/F/G) have been structurally determined to high-resolution by x-ray crystallography or cryoEM ^29–31,36–39^. However, viral infections and virion assembly likely require coordinated interactions among these proteins and the viral membranes, an aspect not directly investigated in many structural studies. Previous studies have characterized the membrane-associated organization of full-length MeV and NDV structural proteins using cryogenic electron tomography (cryoET) ^21,40^. While subnanometer structures of *Paramyxoviridae* matrix protein complexes in association with membranes are not yet available, some studies have revealed that the paramyxoviral M lattice exhibits distinct structural features, different from those of other viruses in the *Mononegavirales* order such as respiratory syncytial virus (RSV) of the *Pneumoviridae* family and influenza of the *Orthomyxoviridae* family ^21,40–42^. For RSV and influenza, these studies reported that the M lattice associated directly with the viral membrane, with their structures reported to have helical symmetry. However, the M lattices of MeV and NDV are predicted to adopt either C2 or C4 symmetry, indicating key differences between these viral families ^21,40,43,44^. Furthermore, subtomogram averaging (STA) of the M lattice in studies of MeV and RSV indicate potential co-assembly of the F glycoprotein with the underlying matrix lattice, although the structural outcomes differed between them. For MeV, individual F trimers appeared to form a regularly spaced, ordered array which appeared to be aligned with some part of the M lattice ^40^. In contrast, RSV F trimers were predominantly arranged as antiparallel pairs that are reported to be in registration with M ^42^. Differences between these studies highlight the importance of understanding the distinct modes of F-M coordination within paramyxoviruses and among related viral families ^28,40,42,44^.

Paramyxoviruses are pleomorphic, with varying sizes and shapes including spheres, filaments, and other forms ^21,40^. In contrast, other members of the Mononegavirales order, such as filoviruses and pneumoviruses, are predominantly filamentous. While the protein-based determinants of filamentous phenotypes are established (helical assemblies of structural proteins), no such model has emerged to describe paramyxovirus pleomorphism. To address this gap in our understanding, we developed a NiV VLP system to systematically interrogate the spatial localization of the four major NiV structural proteins N/M/F/G. This system was leveraged to determine the organization and effect these proteins have on each other and in the context of membranes, in near-native VLPs. Use of cryoET and STA revealed that adjacent M dimers are rotated ∼90° with respect to one another, resulting in the formation of two alternating distinct “holes” in the lattice. We found that F trimers specifically occupy only one of the two holes, and this specificity was lost in the absence of the cytoplasmic tail. G and N were also found to impact NiV VLP pleomorphism. These important structural and morphological findings are important to our understanding of the family *Paramyxoviridae*.

## Results

### Role of Nipah virus structural proteins in VLP formation and morphology

Although paramyxoviral pleomorphism has been observed for over 70 years, the protein complexes or mechanisms responsible for such variability in viral particle morphology remain largely unknown ^45^. To systematically investigate the ability of the NiV structural proteins to form VLPs, we first transiently transfected mammalian expression plasmids encoding each individual structural protein (N, M, F or G) into HEK293T cells (Fig. 1A). Cell lysates and supernatants (cell media fractions containing concentrated VLPs) were quantified by Western blot analysis to determine levels of VLP budding relative to cellular expression, and these samples also were visualized via negative staining transmission electron microscopy (nsTEM) to assess VLP morphology (fig. S1A-C). When visualized by nsTEM, NiV F VLPs and putative M VLPs appeared spherical (figs. S1B top and S1C top) ^32^. Unlike F VLPs which had a clear protein layer decorating their surface, the putative M VLPs had no discernable protein layer (fig. S1B top). These observations are consistent with our prior studies that NiV M and F each are efficient at VLP formation (Fig. 1B and fig. S1A) ^32^.

**Figure 1.**
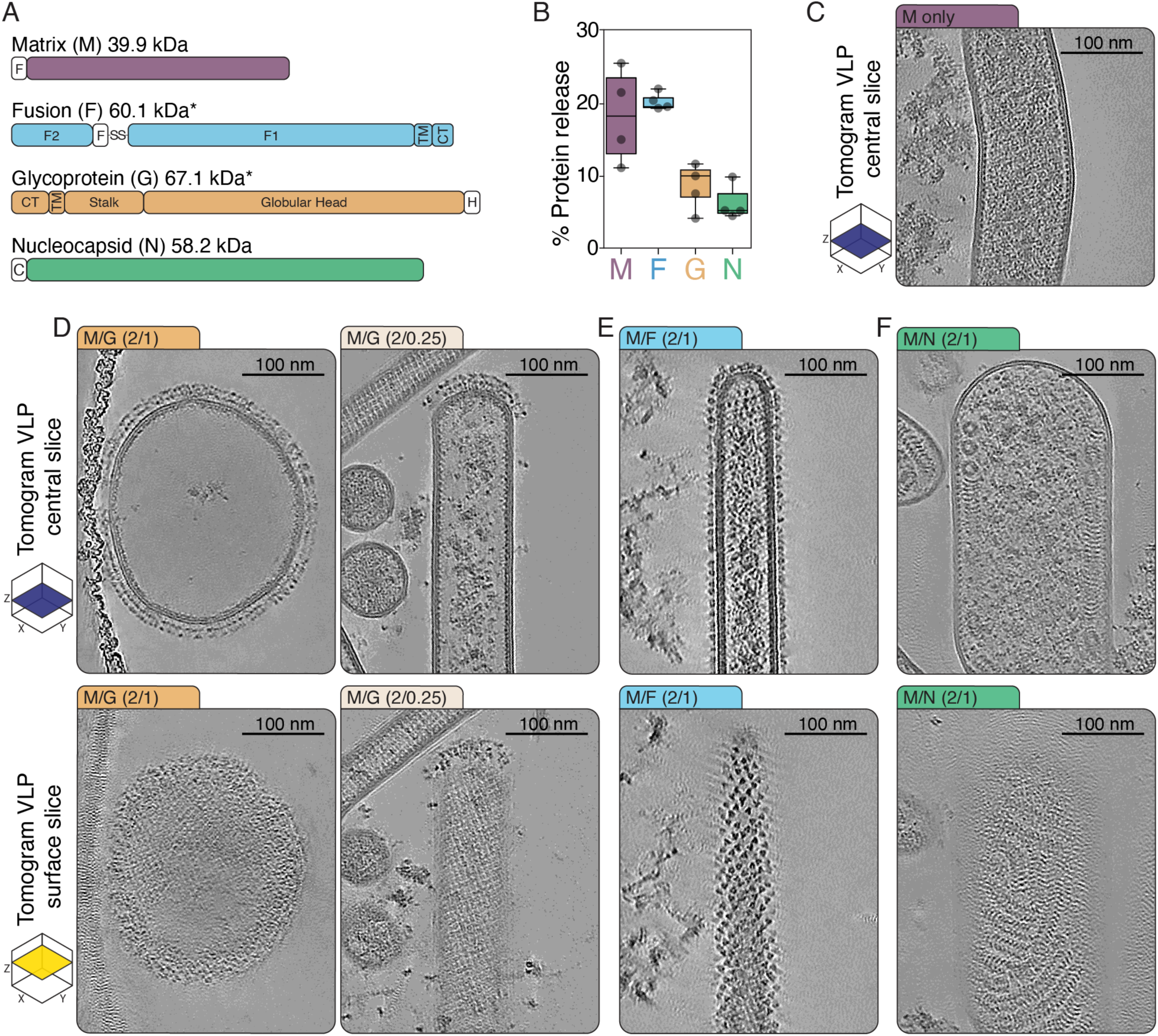
**Role of Nipah virus structural proteins in VLP formation and morphology**(**A)** NiV structural protein schematics and location of tags for the constructs used in this study, F= Flag, H=HA, C=c-myc, TM= transmembrane domain, CT=cytoplasmic tail. **(B)** Relative budding efficiency (% protein released from cells) of each individual NiV protein determined by densitometry of western blots (supernatant divided by cell lysate normalized to total protein). **(C-F)** Cryo-electron tomograms of NiV VLPs; a central or surface Z-Slice (a 10 nm-thick-slice) for several protein combinations displaying the variable morphologies associated with NiV VLPs containing different proteins, ratios represent the plasmid DNA ratio used for transfection.

We predicted that the drying and staining techniques required for nsTEM were altering their appearance in nsTEM. To address this, we vitrified samples for cryoEM in near-native (physiologic salt and pH) buffer conditions to enhance sample preservation, to observe VLP morphology, and to directly visualize the proteins within them in more detail ^46^. Strikingly, when imaged by cryoEM, the NiV M VLPs were mostly tubular, not spherical (fig. S1B bot). By visual morphological analysis the M-only VLPs consisted of 88% tubes, 8% spheres and 4% other morphologies (fig. S1D), with the tubes extending for several microns with an average diameter of 94 nm with a standard deviation of 37 nm (fig. S1D). In contrast, and consistent with nsTEM results, the F-only VLPs were spherical under cryoEM and had an average diameter of 129 nm with a standard deviation of 55 nm (fig. S1C bot and S1E). Thus, in this case for NiV M VLPs, nsTEM does not faithfully capture the native virion morphologies.

To better understand the 3D structure and organization of M VLPs, we collected cryo-tilt-series and reconstructed tomograms. Analysis of reconstructed tomograms revealed electron microscopy density along the inner membrane leaflet of the VLPs, likely the M protein lattice (Fig. 1C) ^21,40,44^. To characterize the protein:protein interactions that may exist between M and the other NiV proteins N/F/G, we co-transfected a consistent concentration of M DNA with increasing concentrations of N/F/G DNA. M DNA was always in excess at 2 ug, and the ratio of M to N/F/G DNA is denoted as 2/(0.25, 0.5 or 1 ug) (Fig. 1D-F and fig. S2A-L). Cell lysates and supernatants of transfected cells were collected, VLPs were purified from supernatants by ultracentrifugation, and protein expression and budding were measured by western blot analysis (fig. S2A-F). Subsequently, concentrated supernatant fractions were prepared for cryoEM and cryoET by vitrification (Fig. 1C-F and fig. S2G-I). Several hundred images and dozens of tomograms were collected for VLPs at every protein combination and ratio, and their morphologies were scored for all M containing VLPs (fig. S2G-L and fig S3 A-B). All samples in this study contained some contaminating vesicles and cellular proteins. For the purposes of analyzing NiV assembly by cryoEM, the term ‘VLPs’ refers exclusively to discrete particles in which the M lattice was visibly detectable.

Co-expression of G with M in co-transfected cells led to an increase in VLP budding as well as a change in the VLP morphology compared to M or G alone (Fig. S2 A, D, G, J). In cryoEM images and tomograms of M:G VLPs, it was possible to directly visualize the full-length G tetramers embedded in the membrane (Fig. 1D and fig. S2G). As the ratio of G to M DNA was raised, more spherical particles were observed (Fig. 1D and fig. S2G), all being extensively decorated with G on membrane surfaces. Interestingly, the M lattice in M:G spheres were primarily discontinuous patches but still appeared to be oligomeric and associated with the membrane (Fig. 1D (bottom)). At low levels of G DNA, when tubes predominated, the small amounts of visible G protein localized primarily on regions of high curvature, i.e. at the tube ends, and at these regions the M lattice was still associated with the membrane (Fig. 1D, fig. S2G). These results indicate that G either induces membrane curvature by directly altering membrane morphology, and/or preferentially localizing to curved membranes.

Like G, the F protein trimer also was visible on the surface of VLPs, appearing as a triangular electron microscopy density (Fig. 1E) ^30,47,48^. In contrast with G, however, F did not promote changes in morphology, with M:F VLPs being primarily tubular, and with F found both at regions of high and low curvature (Fig. 1E and fig. S2 B, E, H, K). Remarkably, we observed that in contrast to the apparently random arrangement of F trimers in F-only VLPs, in M tubes F was regularly arranged, consistent with previous reports for MeV (Fig. 1E and fig. S2H) and suggesting an interaction between NiV F and the NiV M lattice ^40^. At higher concentrations of F the proportion of other types of pleomorphic M:F VLPs increased, suggesting that high densities of F may alter the M lattice (fig. S2H and K).

When assembled into M-VLPs, the NiV N protein formed a characteristic “stacked disc” assembly that was observed both in cryoEM images and in tomograms (Fig. 1F) ^33,38^. The presence of N drastically altered the overall VLP morphology as the VLPs compared to M only VLPs (fig S3 A-B). Unlike the majority of tubular or spherical M:F and M:G VLPs, the majority of M:N VLPs were so pleomorphic that they could not be classified either as spheres or tubes (Fig. 1F and fig. S2 I and L, and S3A-B). Since neither the RNA genome nor the P protein was present in this study, the N polymers likely contain cellular RNA. Although P-independent binding of N to cellular RNA *in vitro* has been reported for Sendai and Measles viruses, our results suggest that N bound to cellular RNA in living cells can still associate with M and that this complex can bud out of cells, suggesting a strong interaction between these two proteins N and M ^17,33,49,50^.

High G or sheering forces can damage large enveloped viruses like orthomyxoviruses, and can result in the tubulation of vesicles ^51^. To address the possibility that the purification procedure or the presence of ectopic tags for tracking the protein of interest affects VLP morphology we performed a key set of control experiments. We generated and transfected new plasmids without protein tags directly into HEK293T cells cultured and adhered to cryoEM grids. The transfected cells were vitrified and imaged by cryoEM and cryoET. Cells were screened at low magnification (3000x) (fig. S4 A,C,E and G), and tilt series near cell edges were collected at high magnification (64000x) (fig. S4 B,D,F, and H). In all four transfections or co-transfections – i.e. M only, M:G, M:F, and M:N – VLPs were observed, and they had the same morphology and protein organization observed for VLPs purified via ultracentrifugation. Therefore, the purification method used here to produce NiV VLPs does not impact protein organization or how the protein complexes impact VLP morphology.

In summary, here we present a robust VLP expression system for producing large quantities of enveloped NiV VLPs from human cells for cryoEM analysis, enabling assembly and structural studies for a BSL-4 pathogen. Notably, these studies yielded several discoveries that critically inform virion assembly and structure for various combinations of the major NiV structural proteins. Comparison of our nsTEM and cryoEM results demonstrate that nsTEM can result in artifacts that alter interpretations. The cryoEM results demonstrate how each NiV protein effects VLP morphology and demonstrates how pleomorphism can occur in paramyxoviruses.

### Structure of the NiV M Lattice

Currently there is no sub-nanometer structure of any paramyxovirus M lattice. Given that these results show a clearly discernable M lattice associated with the VLP membrane, we performed subtomogram averaging. (Fig. 1C-F). VLPs released and purified from cellular co-transfection of M DNA with a low amount of G DNA (M/G, 2/0.25) yielded a sample of sufficient abundance and quality for cryoET data collection, producing 35 tilt series. Analysis of the G conformation on these VLPs was consistent with previous results for four of five paramyxoviruses, and in contrast to reports for Langya Virus (LayV). The G protein on our VLPs predominantly appeared in the “two-heads-up and two-heads-down” conformation as a tetramer, and the heads appeared to display high rotational freedom (fig. S5D middle) ^29,37,52–55^. As shown for M:G VLPs in figure 1, tubes in this data set contained little or no G in the VLP membrane along the sides of the tubes where the M lattice was most clearly observed (fig. S5A). Tube diameters varied from 50 to 100 nm, consistent with the M-only VLPs (Fig.1C and fig. S1D and S5A).

To determine the structure of the M lattice, subtomogram averaging was performed on particles extracted from the reconstructed tomograms (fig S5B-C and S6A-D). Consistent with previous results for other paramyxoviruses ^21,40,56,57^, our ab-initio reconstruction and subsequent alignment revealed a checkerboard array of M-dimers separated by holes, lining the inner membrane leaflet (fig. S6E). Further subtomogram averaging and 3D-classification revealed that M dimers opposite one another across the hole were rotated by 90 degrees. This resulted in two distinct classes of M-dimers that, when projected into real space, were alternating in both Alternating along and around the spline of the tube at the membrane (Fig. S6F). Subtomograms were extracted into a larger box size to account for the two M-dimer orientations (9 dimers total) and subjected to further refinement. The resulting map led to the following main conclusions: First, adjacent M dimers are related to each other by a ∼90° rotation, with the dimer:dimer interface requiring interactions between three monomers instead of two as previously hypothesized ^40^. Second, as a result of this rotation and the approximately rectangular dimensions of the M dimer, the resulting lattice contains two distinct holes (herein termed A and B) that differ in size, shape and electrostatic properties (fig S7A-B). Finally, the M lattice is formed through a single dimer:dimer interface, in contrast to earlier models that proposed two distinct dimer interfaces ^40^.

The final subtomogram average of 9 dimers had an estimated resolution of 7Å allowing for modeling (Fig. 2A and fig S6G). This structure allows for the visualization of the interface between two M dimers, as well as the two distinct holes they form (Fig 2B-C). A *membrane proximal* orthoslice reveals that hole A is smaller next to the membrane (4.4 nm across) while hole B is larger (6nm across) (Fig. 2B and 2C). In contrast, a *membrane distal* orthoslice reveals that the relative sizes of these holes are reversed, with hole A being larger (7.8 nm) away from the membrane, and hole B being smaller (4.4 nm) (Fig. 2B and 2C). The two tetramers of dimers show no obvious differences in hydrophobicity, but hole A is more positively charged than hole B (fig. S7A-B). Structural comparisons between this model with a previously published crystal structure are consistent with the model in which the NiV M dimer complexes with PI(4,5)P_2,_ as we identified several key structural motifs in our map and model that more closely resemble the holo form (fig. S7C-D) ^39^. These findings describe the first sub-nanometer structure of a paramyxovirus M lattice, revealing a novel arrangement of M proteins that have implications for glycoprotein arrangement.

**Figure 2.**
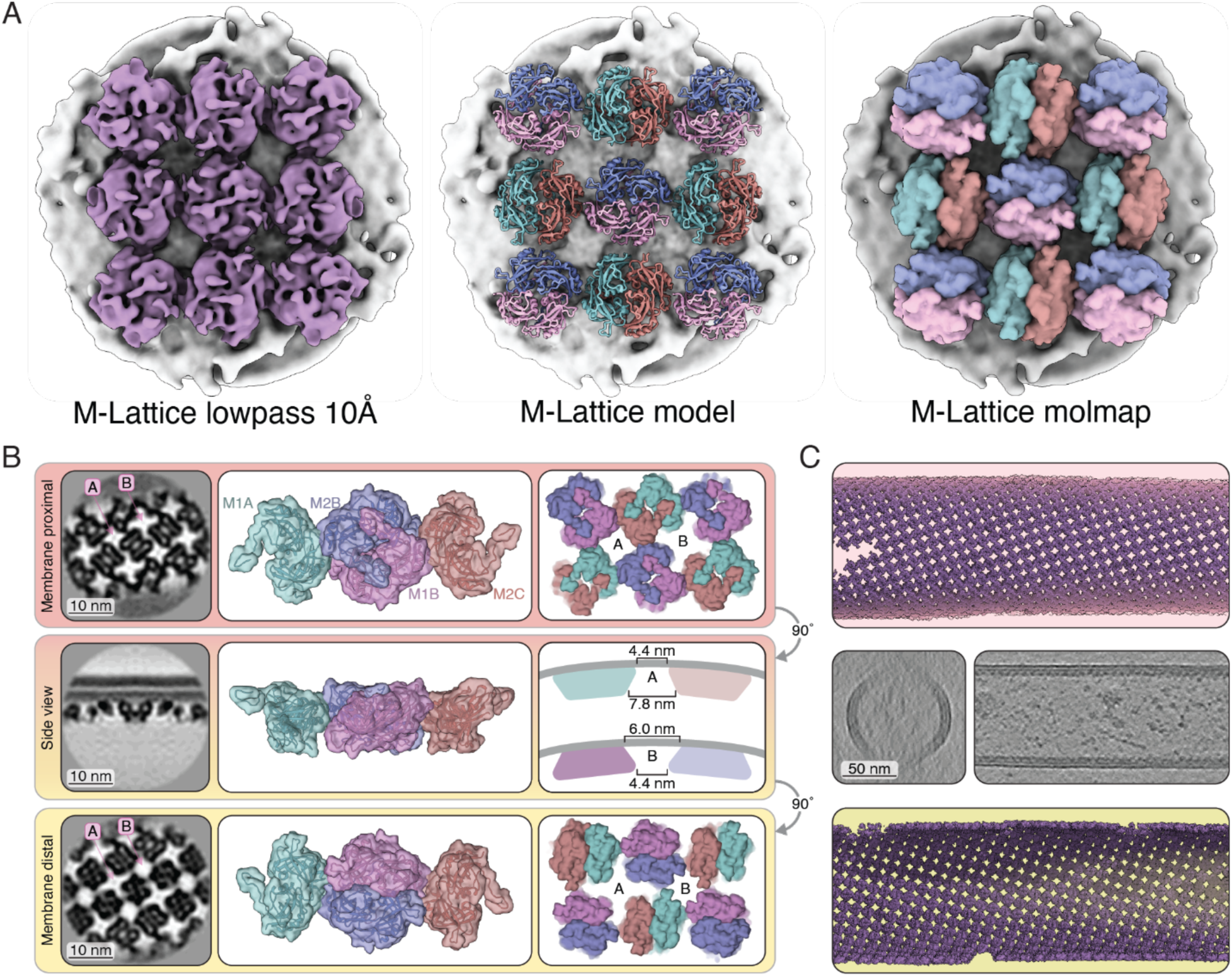
Subtomogram average of the NiV M lattice. **(A)** Lowpass filtered electron density map (left), molecular model (middle) and corresponding molmap (molmap=map-like density generated from a model at a resolution of 8Å) of the NiV M Lattice (**B)**, Descriptions of several orthoslices and orientations of the NiV M oligomeric assembly in three viewing angles. Scale bars 10 nm (**C**), Alternative viewing angles of NiV M oligomeric assembly generated by lattice mapping.

### Subtomogram average of NiV F and M:F complexes

As detailed above, F-only VLPs were predominantly spherical while M:F VLPs were predominantly tubular (Fig 1 and fig. S1C, 1E and S2H, S2K). The triangular F protein in F-only VLPs appeared to be randomly distributed on the VLP surface (Fig. 3A). To investigate this observation, we conducted subtomogram averaging of F proteins on F-only VLPs, resulting in a map with an estimated resolution of ∼9Å (Fig 3B and fig. S8A-F). Docking of a soluble F trimer model (missing it’s transmembrane and cytoplasmic domains—PDB:8DNG) into our map revealed that the overall shape of the ectodomain of the full length NiV F is consistent with commonly studied soluble versions (Fig 3B and fig. S9A) ^47^. Importantly, projection of the refined F protein positions from VLPs into real space shows that F is randomly distributed on the surface of F-only VLPs (Fig 3C).

**Figure 3.**
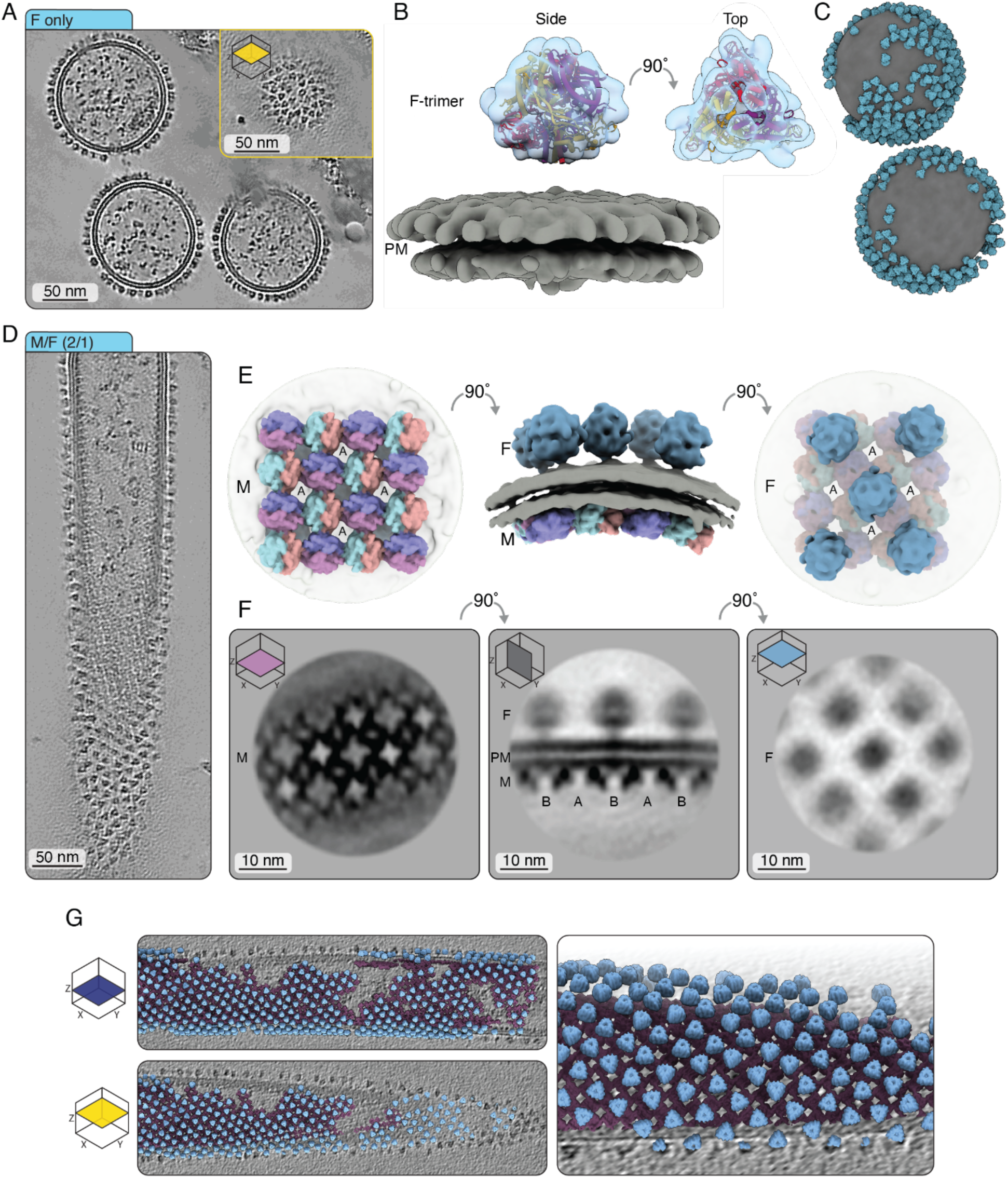
Subtomogram average of full-length NiV F and M:F Complex. **(A)** Cryo-electron tomogram of NiV F-only VLPs; a Z-Slice (10 nm). **(B)** Subtomogram average of full-length NiV F (blue) in complex with human cell membrane (gray) docked with PDB 8DNG. **(C)** Rendering of the tomogram in (A) displaying the real-space position and angle of each F trimer (particle positions and poses extracted from particle tables used to generate the subtomogram average). **(D)** Cryo-electron tomogram of NiV M:F VLP; a Z-Slice (10 nm) **(E)** Isosurface model of MF Complex, F and membrane shown as maps lowpass filtered to 10Å, M Lattice displayed as a molmap model of 10Å (molmap=map-like density generated from a model at a resolution of 10Å). **(F)** Orthoslices from M:F subtomogram average with annotation of holes in M lattice. **(G)** F and M maps (lowpass filters of 15Å) reprojected back in 3D space using particle poses used to generate M:F subtomogram averages with central coordinates for F and M respectively.

In contrast to F-VLPs, M:F VLPs were predominantly tubular, and F trimers in M:F tubes formed regularly arranged rows on the tube surfaces (Fig. 3D). Therefore, we hypothesize that the cytoplasmic tails of F proteins might assemble with the M-lattice, resulting in the observed regular arrangement. Subtomogram averaging of the M:F complex was performed, resulting in an electron microscopy density map of the M lattice with an estimated resolution of 9Å (fig. S10A-D). Subsequently, subtomograms were re-extracted with a box size that included both the M and F layers (fig. S10E). Without changing the subtomogram position or angle, map density corresponding to the F-trimer was present above only hole B of the M lattice (Fig. 3E-G and fig. S10A-E). At higher map thresholds, we observed a map density protruding from the inner membrane leaflet directly below the F trimer. Despite significant effort, and the tantalizing nature of this observation, we were unable to assign the observed map density as part of the F cytoplasmic tail (fig. S9E). However, the complete localization of the F-trimer to hole B in the M lattice strongly suggests an underlying mechanism.

Subtomogram averaging of the F trimer on M:F VLPs did not lead to an improvement in the estimated resolution compared to the F map obtained from F-only VLPs. However, analysis of the aligned F subtomogram positions and orientation relative to the membrane surface allowed us to compare the spatial and angular distributions of F on F-only spheres from the distributions of F on M:F tubes. The angular freedom of F trimers on spheres was ∼5° and on tubes ∼10° (Fig S9B and S9D). Also, the spacing between F trimers on F-only spheres ranged from 5-15 nm and were randomly distributed, whereas the spacing of F trimers on M:F tubes clustered around 12 nm and were ordered (fig. S9C and S9E). Interestingly, we could not resolve robust stalk density for any F protein structure in F only or M:F VLPs, likely due to size, flexibility or issues regarding CTF correction near the membrane.

Given the above results, we hypothesized that the F cytoplasmic tail determines the regular distribution of F on the surface of M tubes. To test this hypothesis, we engineered an F protein (F(trunc)) lacking most of the C-terminal cytoplasmic tail except the KKR motif that is essential for proper trafficking to the plasma membrane (fig. S11 A) ^58^. Tomograms of M:F(trunc) VLPs had no regular spacing of the F(trunc) trimers demonstrating a loss of the regular structural organization observed previously in wild-type F protein trimers assembled on M tubes (fig. S11 B). STA analysis of the M lattice in M:F(trunc) VLPs resulted in a map that was indistinguishable from the M lattice from M only or MF VLPs (fig. S11 C-E). In contrast with M:F VLPs, for M:F(trunc) VLPs there was no density corresponding to F further demonstrating that F(trunc) trimers had no regular organization. (fig. S11 C and E).

### CryoET of NMFG VLPs

To study VLPs that more closely resemble whole NiV virions, we co-transfected N/M/F/G, into cells and assessed the released VLPs. While the ratio of NiV proteins in infectious particles is not known, members of its order *Mononegavirales,* exhibit a transcriptional gradient from the 3’ to 5’ end of the genome, with gene location influencing expression levels ^22^. Extrapolating from this, the likely expression hierarchy is N > M > F > G. To approximate these relative expression levels, we chose DNA ratios of 12:9:3:1 for N, M, F, and G, respectively.

N/M/F/G VLPs in tomograms were highly variable in both their protein content and morphologies. Some VLPs had clearly defined densities for all four viral proteins (Fig. 4A), and the localization of viral glycoproteins was consistent with the data for M:F and M:G VLPs. Trimers of F were ordered on flat edges of the VLP surface, while in the same particle, G protein tetramers were localized to regions of high curvature (Fig. 4A). To quantify these observations, central Z-slices of 24 N/M/F/G VLPs were analyzed in regions of high and low curvature (fig. S12A). F trimers were observed on areas of both high and low curvature at 4.86 and 5.96 F trimers per 100 nm of membrane length, respectivel*y* (fig. S12B). By contrast, G tetramers displayed approximately a 10-fold stronger preference for regions of high curvature, with 3.64 G tetramers observed per 100 nm on regions of high curvature but only 0.36 G tetramers observed per 100 nm on regions of low curvature (fig. S12A-B). Using this data, an interactive structural model was constructed which provides a glimpse into Nipah virus structural protein organization in particles (Fig. 4B Interactive Tour).

**Figure 4.**
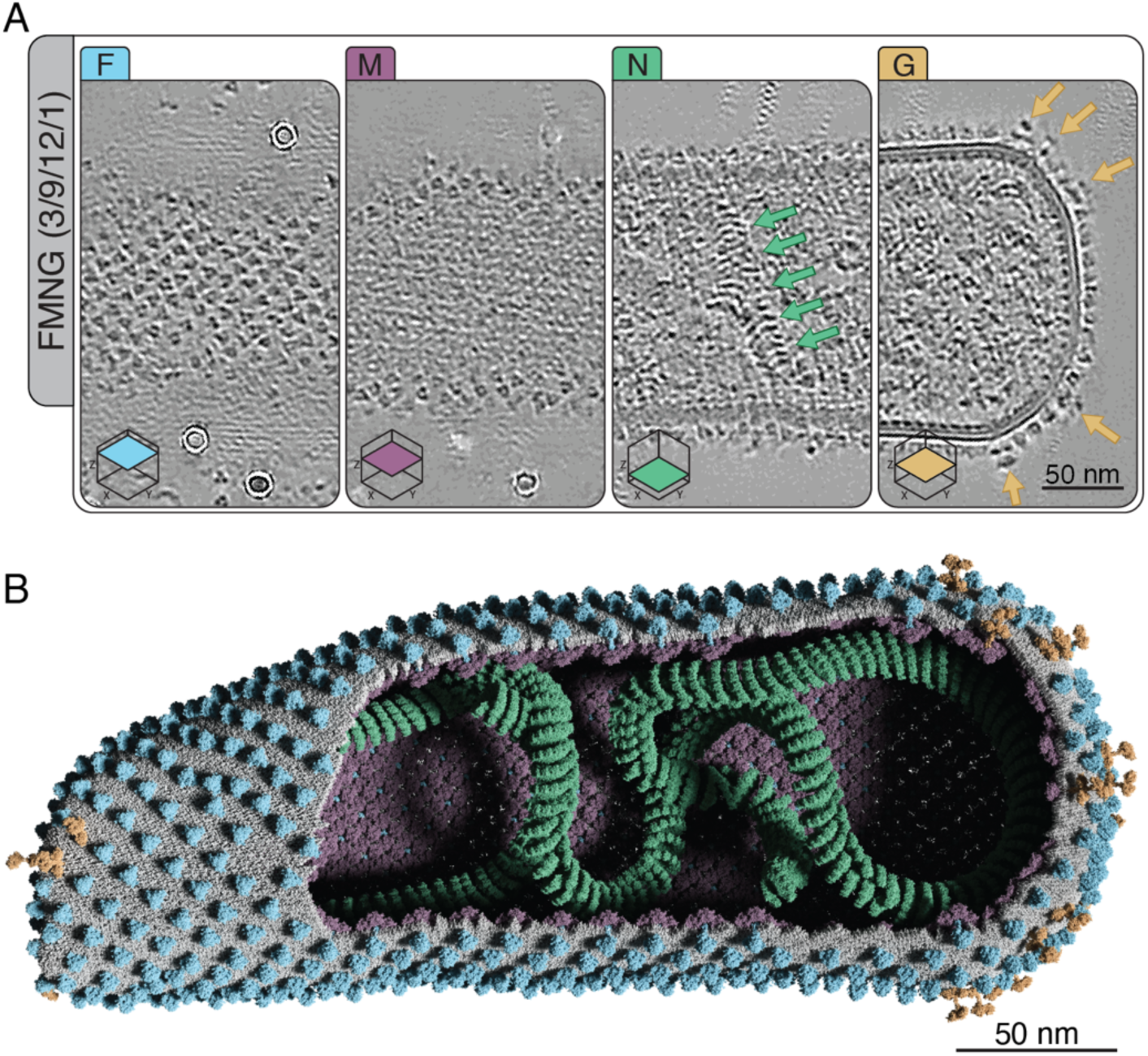
CryoET and Model of NMFG VLPs. **(A)** A tomogram of an NMFG VLP highlighting the location and organization of structural proteins. **(B)** Idealized 3D model built from tomogram in (A), rendered in Blender using the MolecularNode addon. This model was generated to enhance interactive scientific communication, and is publicly available <u>online</u>.

## Discussion

Using cryoEM techniques to analyze VLPs assembled with combinations of co-expressed viral structural proteins, we have characterized the architecture of the paramyxovirus NiV. An advantage of this expression system is that for each structural protein, the spatial compartmentalization and influence on morphology can be assessed independently. The matrix protein, M, is sufficient to bud tubular particles from cells. The structure of the NiV M (paramyxovirus) lattice presented here is unique, with interactions not previously described for M lattices of the RSV (pneumovirus), Ebola (filovirus) and Influenza (orthomyxovirues). The NiV lattice consists of two M dimers, rotated approximately 90 degrees with respect to one another, and continued propagation of this unit results in a lattice with two distinct overlapping “tetramers of dimers”, resulting in two distinct “holes” designated (A and B). Although the fusion glycoprotein F can bud VLPs with random distribution of F-trimers on the membrane, in the presence of M, the F trimer becomes ordered and localizes above only the B-hole. We confirmed via a combination of mutagenesis and subtomogram averaging that the short cytoplasmic tail of F is responsible for the preferential sorting of F trimers into hole B. Expression of M and G results in tubular and spherical VLPs where G is predominantly localized to highly curved parts of the membrane, such as the ends of tubes. Co-expression of N with M results in irregularly shaped VLPs containing N oligomers (presumably assembled with cellular RNA).

While the co-transfection system implemented here has many advantages for uncovering the roles of different structural proteins in virus assembly, particularly with high-containment viruses, limitations in interpreting the relevance to a real NiV infection need to be considered. We eliminated the possibility that epitope tags or ultracentrifugation significantly affected our results regarding the oligomerization of M, spatial compartmentalization of F and G, and the ability for N to drive particle pleomorphism, since all were still observed in VLPs captured without ultracentrifugation. However, an inherent limitation for all transfection experiments is that the viral proteins were expressed at high levels, a requirement for generating sufficient quantities of VLPs for cryoEM analysis. It is unknown how the expression levels of these proteins in transfected cells compare with levels in authentic NiV infected cells. Finally, the effects of viral proteins other than N, M, F, or G on virion assembly and structure are unknown and would require further studies investigating structures of live NiV in a BSL4 environment.

The matrix protein of all known paramyxoviruses and many other Mononegavirales members are reported to be predominantly dimeric ^15,21,42,57^. However, M proteins differ greatly in sequence and length, and so it is reasonable to predict that they do not have the same lattice arrangements between families ^21,40,56,59^. To date, the highest-resolution matrix lattice structure is from the pneumovirus RSV. RSV M differs significantly in sequence and length (256 residues for RSV, 352 residues for NiV) and lattice symmetry (RSV, helical; NiV, helical in tubes but C2/pseudo-C2 locally). Additionally, the spacing and geometry between adjacent dimers vary across published M lattices. In viruses with helical arrays like Rhabdoviruses, Orthomyxoviruses and Pneumoviruses, a single dimer constitutes the asymmetric unit ^41,42,60^. Notably, for NiV the asymmetric unit contains two M dimers, suggesting that in C2 or pseudo-C2 symmetric lattices, matrix proteins may function as dimer pairs, that can assemble as a helix but can also assemble as “sheets” of M lattice. The spacing between matrix protein dimers in the lattice may also contribute to the apparent fragility of the lattice. In many instances, we observed lattice disruption at sites of curvature or bending. Similar structural fragility has been reported for the pseudo C2-symmetric VP40 lattices of Ebola virus, in contrast to the more stable helical matrix arrays observed in rhabdoviruses which do not display high pleomorphism ^61^. Mechanistically this phenomenon is likely due to the increased surface area of dimer:dimer contacts present in helical arrays and to the presence of the large gaps indicative of a non-helical C2-symmetric lattices. Given the high structural homology of paramyxovirus M dimers we predict this lattice structure and dimer rotation might be observed in other paramyxoviruses. These observations suggest that lattice symmetry may influence structural stability.

Our model further refines this concept, demonstrating that the NiV matrix lattice is formed through a single interface between two rotated M dimers, propagated out continuously over the lattice structure. This single repeating interface, which leads to the formation of the two types of holes in the lattice, provides a simplified model compared with previous models that proposed two distinct interfaces (monomer:monomer and dimer:dimer) for lattice formation ^21,40^. Our model is further supported by observations that F localizes specifically to one of the lattice holes, and that distinct charge distributions are present within holes formed by four M protein dimers. While we have proposed a structural/functional role for one lattice hole, the function of the other hole type remains unresolved. Perhaps the other hole is involved in accommodating host cellular proteins, a hypothesis that warrants further investigation. It is noteworthy that the conformation of the M dimer observed in our study aligns with the previously crystallized holo form of NiV M bound to phosphatidylinositol 4,5-bisphosphate (PI(4,5)P₂), further supporting the role of this lipid in viral assembly ^39^

While an MF interaction has been suggested for the paramyxovirus MeV ^40^, the regular ordering of NiV F that we observed on the M lattice is the first such visualization of its type for henipaviruses. Our results indicate that the interactions between the NiV F cytoplasmic tail with hole B of the M lattice are indeed specific but are likely weak and/or transient, and/or that the F cytoplasmic tail lacks significant secondary structure. Our study is also the first to show that the F cytoplasmic tail is required for F trimer arrangement, and assign this organization to a specific structural motif in the M lattice (hole B). We speculate that hole B, which possesses a larger membrane-proximal space, is sterically preferred to better accommodate the cytoplasmic tails of F. Rigid-body docking of full-length Alphafold3 F models into our M:F map suggest that the dimensions of F could allow for cytoplasmic tail interactions between both membrane distal and membrane proximal matrix lattice surfaces. Our observations and those for Measles, two separate viral genera, suggest that a regular arrangement of F may be a characteristic of the *Paramyxoviridae* family. High expression levels of F led to more pleiomorphic VLPs, with FM tubes containing more bends. We hypothesize that introduction of the F cytoplasmic tail domains between M dimers destabilizes the M lattice, which increases its flexibility, resulting in increased VLP pleomorphism. An intriguing idea is that this may be advantageous for membrane fusion and viral entry. Previous reports have suggested that F in VLPs may form hexamers of trimers or a dimer of trimers; however, we did not observe this arrangement in any VLP sample, suggesting this may be an artifact of soluble ectodomain studies ^36,62,63^.

Neutralizing antibodies often target the apex of the paramyxovirus F proteins ^30,47,64,65^. Since F trimers on the F-only VLPs studied here are densely packed on the virion surface, we propose that neutralizing antibodies targeting the F trimer apex could be sequestered by non-infectious F-VLPs, which would support their role as immunological decoys *in vivo,* an idea proposed previously ^19,30,47,58,65,66^. Conversely, due to the wider spacing of F trimers in the context of M lattice, additional epitopes might be accessible. Thus, future vaccines utilizing M:F VLPs might help direct the immune system towards novel epitopes. A recent study has identified antibodies that bind on the lateral and basal surfaces of NiV F ^47^. Given our findings it is possible that such antibodies are more effective against infectious particles.

At least 3 of the four NiV structural proteins studied here play a role in VLP morphology. Pleomorphism appears to be determined by the relative concentrations of the different viral structural proteins present in any given particle. For example, while M itself forms tubes, high levels of G or N expression in presence of M lead to spherical or irregular shaped VLPs. Given its position in the genome, it is likely that G expression during infection is far lower than what was represented here, and that the high G:M spheres may not be physiologically relevant. Thus, we predict N:M interactions would drive the majority of pleomorphism observed during live viral infection. However, in particles which contain a low amount of G, which may be physiologically relevant, G tetramers preferentially localized to regions of high curvature. Because viral–membrane fusion begins with G binding to the ephrin B2 receptor ^67^, we propose that G locally modulates the M lattice and/or membrane biophysical properties, a possible function that may be required for fusion. The F trimers near patches of G would then be responsible for initiating membrane fusion due to the role of G in F-triggering but the role of F trimers distant from G patches remains unclear ^68–70^.

It is reasonable to predict that infectious NiV particles, containing all structural proteins and a complete genome, would be pleomorphic. For influenza virus, both spherical and filamentous virus particles are observed, with the latter being more resistant to neutralization ^71,72^. For NiV a proportion of the virus particles might also be filamentous, lacking nucleocapsid assemblies but exhibiting areas rich in viral glycoproteins, and such non-infectious VLPs could serve as decoys for neutralizing antibodies ^73^.

Previous studies of NiV and MeV postulated that the relative concentration of M at the surface of an infected cell might determine whether budding viruses are spherical or tubular. ^56^ Although we cannot rule out changes due to expression levels, we predominantly observed non-tubular M particles only in the presence of high concentrations of G or N. The phosphoprotein (P) for paramyxoviruses is reported to sequester free N protein when the latter is not bound to the viral genome ^17,33,49,50^. However, recent cryoET studies of Mumps virus (MuV)-infected cells showed that the vast majority of N in the cell is polymerized into highly variable lengths of nucleoprotein complexes; although it is unknown if the observed N polymers encapsulate viral or host RNA ^74^. How NiV P expression affects virus morphology remains to be assessed directly. Furthermore, it can be speculated that N polymers assembled with the viral genome could induce some constraints on virion size and affect pleomorphism differently than what we observed here. Future studies with infectious NiV particles will be required to decipher this possibility.

Taken together, our results describe the first sub-nanometer cryoEM structure of a paramyxovirus M protein lattice, as well as the placement of G, F, and N proteins associated with that lattice, thus providing structural insights into the complex and dynamic NiV protein:protein interactions leading to pleomorphism. The results are summarized in an interactive tour to provide support for scientific communication (Interactive Tour). The functional consequences of pleomorphism for NiV and for other paramyxoviruses remain to be explored. Recent studies with HIV-1 have revealed the capsid to be a potent antiviral target for pharmaceuticals ^75^. Perhaps such drugs also could be developed for paramyxoviruses such as NiV.

## Acknowledgments

The authors thank Volker M. Vogt for discussions and help with writing; William Wan for discussions regarding data reporting, validation and Matlab Code; Mariena Silvestry Ramos, Savannah G. Brancato and Katie Spoth for help with microscope maintenance and data collection; the staff and facilities at the Cornell Center for Materials Research (CCMR) and the Electron Imaging Center for Nanosystems (EICN) at the University of California, Los Angeles’s California for NanoSystems Institute (CNSI) for their support and maintenance of the microscopes.

## Author contributions

VVU: Designed protein constructs, generated and purified VLPs, prepared TEM grids, analyzed and assessed VLP morphology, collected all negative stain and cryoEM data, processed and solved structures, analyzed all data present in the manuscript, prepared figures, and wrote the manuscript. JFL: Generated and purified VLPs, analyzed and assessed VLP morphology, prepared figures. NE: Generated and purified VLPs, analyzed and assessed VLP morphology. CR: wrote scripts for subtomogram averaging. ACH: Generated and purified VLPs, analyzed and assessed VLP morphology. MO: Collected cryoET data and wrote scripts for subtomogram averaging. LA: Built molecular model in meoscape. FKMS: Provided input for analysis and interpretation, collected cryoET data, and wrote scripts for subtomogram averaging. HCA: Provided input for analysis and interpretation of results, supervision, funding, manuscript writing. RAD: Processed and solved structures, prepared figures, analyzed and interpreted results, provided supervision and funding, helped with manuscript writing.

## Data and materials availability

The structural Model for NiV M Lattice is deposited into the protein data bank (PDB, https://rscb.org/). The PDB ID is 9NQY. The subtomogram average map for the NiV M Lattice is deposited in EMD-49696.

## Materials and Methods

### Plasmids

Codon-optimized sequences for NiV N, F, F(mut) and G were inserted into pcDNA3.1 expression vectors and tagged with c-myc, 3X-FLAG, and HA tags, respectively. The F and G plasmids were published previously ^19,32,76,77^. Codon-optimized NiV M was inserted into a pCMV-3X-FLAG vector with an N-terminal FLAG tag as described previously ^78^. All untagged plasmids used for cellular tomography and control experiments were the exact same coding sequence and plasmid backbone as their tagged counterparts.

### Cell Culture and Viral-like Particle (VLP) production

HEK293T cells were maintained in Dulbecco’s Modified Eagle Medium with 10% Bovine Calf Serum and 1% Penicillin/Streptomycin, and grown at 37°C with 5% CO_2._ Tissue culture plates at ∼80% confluency were transfected with either 3 µg/well (6-well plates) or 30 µg (15cm plates) of DNA and 12 µg (6-well plates) or 120 µg (15cm plates) of polyethylenimine (PEI) in Opti-MEM for 4 hours. Transfections were allowed to continue for 48 hours. Cells were lysed in RIPA buffer with c0mplete protease inhibitor (Sigma) for cellular protein analysis and the media were harvested for budding analysis. Media were loaded on a 20% sucrose cushion and ultracentrifuged for 90min at 110,000xg. Concentrated virus preparation was then loaded onto a 3-52% OptiPrep (iodixanol) discontinuous gradient and spun at 200,000xg for 14 hours in a SW60ti swinging bucket rotor. Fractions were taken and assessed for presence of VLPs via western blot analysis. Fractions containing viral proteins of interest were pooled and diluted with 50mM Tris pH 8.0, 150mM NaCl and 1mM EDTA to reduce the concentration of Optiprep. VLPs were then pelleted and resuspended in the same buffer. Samples were stored at 4°C until cryo-preservation.

### Western Blotting

Western blot analysis and densitometry were conducted as previously described ^79^. Briefly, SDS-PAGE was conducted and proteins transferred to a PVDF 0.45um pore size membrane. Blot was blocked in Intercept Blocking Buffer (LI-COR) followed by staining with primary mouse or rabbit antibodies against epitope tags overnight. Primary antibodies rabbit polyclonal anti-HA (Sigma Aldrich Cat#H6908), mouse monoclonal anti-FLAG (M2 Sigma Aldrich Cat#F1804), and mouse monoclonal anti-Myc (Invitrogen Cat#MA-21316) were used at a 1:1000 dilution overnight at 4°C. Secondary antibodies were goat anti-rabbit or mouse conjugated to Alexa Fluor 488 or 647 (Invitrogen Cat#A11001, A21236, A11008, A21244) at a 1:2000 dilution for 1 hour at 22°C. Blots were thoroughly washed with Phosphate-Buffered Saline (PBS) with 0.2% v/v Tween 20 prior to exposure and imaging in a Bio-Rad Gel Imager. Densitometry analysis was conducted using Image Lab (v6.0.1).

### CryoEM sample preparation

NiV VLPs were diluted with 10 nm gold fiducials coated in BSA. Quantifoil 300 mesh Au R2/2 were glow discharged at 20mA for 45s. 3uL of sample was applied, blotted for 3-4s and plunge frozen into liquid ethane using a Leica GP2 grid plunger (Leica Microsystems).

### CryoEM data acquisition

CryoEM images for morphological analysis are detailed below in Table S2. Images were acquired on a Talos Arctica cryo-electron microscope (TEM, ThermoFisher Scientific) operating at 200keV using SerialEM (v4.0.6) ^80^. Data was collected using a Gatan K3 direct detector equipped with a BioQuantum energy filer operated in counting mode using a slit width of 20eV. VLPs were collected with a total dose of 50e^-^over 50 frames.

### CryoEM image processing and scoring of VLP morphology

Movies of VLPs were imported into Relion (v4.0) for preprocessing ^81^. Movies were gain-normalized, frames were aligned with MotionCor2 and CTF was estimated with CTFFind (v4.1) ^82,83^. The manual picking tool within inside Relion 4 was used to visualize and score the morphology and particle contents of VLPs.

### CryoET data acquisition

Tilt series for visual analysis and sub tomogram averaging were acquired on either a Talos Arctica or Titan Krios cryo-electron microscope (ThermoFisher Scientific) operated at either 200keV or 300keV respectively (Table S1 and S2). Data was collected on both microscopes using a Gatan K3 direct detector operated in counting mode equipped with a BioQuantum energy filter utilizing a slit width of 20eV. The nominal magnification for tilt series acquisition on the Talos Arctica and Titan Krios were 63,000x and 64,000x corresponding to a calibrated pixel size of 1.33 and 1.39 Å/pixel respectively. Tilt images were acquired using a dose-symmetric tilt scheme (Hagen scheme) from +60° to-60° at 3° increments starting at 0° and reversing direction every 2 tilts.^84^ A total dose of 143.5 (300kV) and 150e^-^/Å^2^ (200kV) was used over the entire tilt series (41 tilts) where each tilt received ∼3.5 and ∼3.65e^-^/Å^2-^ over 8 or 10 frames respectively.

Tilt series were collected over a range of defocus between-1 and-4 microns in steps of 0.5 microns. Specific microscope and data acquisition values are provided in Table S1 and S2.

### Tilt-Series preprocessing and tomogram reconstruction

SerialEM (v4.0.6) was utilized to collect binned K3 movies (5760×4092) for each tilt, frames were gain-normalized and aligned on-the-fly using a SerialEM implementation of MotionCor. Microscope collection resulted in ordered stack files containing all tilts. Tilt series were visualized in 3dmod where bad tilts (images where the particle of interest was occluded or lost during collection) were removed and remaining movies were dose filtered to down weight movies with high exposure using TOMOMAN ^85,86^

Cleaned and dose filtered tilt series were cropped and CTF estimation was performed using CTFFind4 (v4.1.1) on each tilt individually ^82^. Tilt series were then aligned through manually seeding and tracking of gold fiducials and then reconstructed with VLPs of interest centered within the volume using IMOD ^86^.

Final tomograms for subtomogram averaging were generated using the ctfphaseflip function in IMOD resulting in CTF-corrected unbinned and binned tomograms for averaging in Dynamo^86,87^.

Final tomograms for morphology analysis and depiction were generated through the TOMOMAN wrapper of cryoCARE ^85,88,89^. 10 frame movies for each tilt were split in half using the TOMOMAN wrapper of MotionCor2. Odd/Even tilt-series were then cleaned, dose-filtered, and aligned using TOMOMAN ^85^. CTF was determined using the TOMOMAN implementation of tiltctf ^90^. Odd/Even ctf corrected tomograms were then reconstructed using IMOD and gold was erased. Odd/Even tomograms were used to train the cryoCARE neural network for 100 epochs with a learning rate of 0.0004.

### Subtomogram averaging

#### NiV M Lattice

The complete subtomogram averaging pipeline for the NiV M lattice is detailed in fig. S6. Since the NiV M VLPs are tubes of a roughly consistent diameter, we were able to use a geometric model to define initial positions and Euler angles using Dynamo/TOM/AV3 utilities^91^. For all tomograms, a spline running down the center of the tube was modelled in 3dmod and the radius was measured. A single tilt series collected at high defocus was reconstructed and binned by 8 for initial model generation. Positions corresponding to M lattice and membrane were extracted in Dynamo to generate an ab-initio model to serve as a model to align the full data set ^87^. Two sets of alignments were conducted, first a large Z shift was applied to align the membrane for all subtomograms, followed by translational and angular alignments to align the M lattice. Angular refinement in Dynamo was conducted until a strong initial model was generated.

In total, 24 tomograms corresponding to 26 tubes were utilized for the final reconstruction with the full processing pipeline outlined in a supplemental figure (fig. S6A-D). Subtomogram alignment of the entire data set began by extracting and aligning all bin8 subtomograms against the ab-initio model in Dynamo ^87^. Since we oversampled the initial positions, Dynamo functions were used to remove overlapping and duplicate particles. Particles that failed to align to the reference were also removed based on their lower cross correlation.

Particles were subjected to several rounds of alignment and cleaning followed by unbinning from bin8 to bin2. At bin2 the data set was imported to Relion 4 tomo and subjected to 3D-classification and alignments ^81,92,93^. At this point we observed a split in the data set that resulted in two different classes of particles corresponding to the two orientations of the M lattice. The asymmetric unit of the lattice consisted of two dimers, rotated approximately 90° with respect to one another. We then chose to apply C2 symmetry for all further processing. Both classes were processed separately in Relion 4 Tomo for several rounds of pose refinement, frame alignment and per particle CTF-determination. Finally, the box was shifted onto the central dimer and refined to best depict the two tetramers of dimers and yield the final map reconstruction (fig. S6G).

#### F on F VLPs

Subtomogram averaging for full-length NiV F on the surface of VLPs was conducted as described for the NiV M lattice with some minor differences. The complete pipeline is described in fig. S8. Unlike M Tubes, F spheres were not uniformly shaped so instead of geometric based picking we elected to manually annotate the membrane of the F spheres in Dynamo ^87^. A gaussian filter of 2.76Å or lowpass filter of 10Å was applied for visualization purposes.

#### M:F Complex

Subtomogram averaging for the M:F complex was conducted as described for the NiV M lattice with some minor differences. The complete pipeline is described in fig. S10. Since F occupies only one of two tetramers of dimers, 3-D classification in Relion tomo was unnecessary and the classes were split into hole A and hole B automatically.

#### M:F(trunc) Complex

Subtomogram averaging for the M:F(mut) complex was conducted as described for the NiV M lattice with some minor differences. Tomograms used for subtomogram averaging in Dynamo were reconstructed with novaCTF instead of ctfphaseflip and the initial 2-D CTF estimation was calculated with tiltctf ^90^.

#### Positional and Angular Distribution of F ectodomain

The angular distribution of NiV F ectodomains in the context of F only spheres or M:F tubes was determined by first extracting the Euler angles from the particle tables used for subtomogram averaging of the F trimer. Psi and theta angles were converted to a position on a unit sphere (phi = 1) then converted to polar angles. The angular difference between the aligned position and the reference positions were calculated and converted back to degrees. The reference position and angle for the F spheres was defined by manually annotating the membrane and the center of the VLP wherein the initial Euler angle was defined by a vector from the center of the VLP through the position on the membrane. For M:F, the reference position was defined as the center coordinate and angle of the M lattice below the F protein. Differences between the aligned angle and initial angle were plotted as pie chart and displayed as a slice of that pie chart in fig. S9B. The distance between F trimers on F-only spheres and M:F tubes was determined in both 1D and 3D. For the 1-dimensional analysis, we used the Subset selection function inside of Relion 4 Tomo to remove duplicates of a given inter-particle distance. We calculated the number of particles removed in steps from 5Å up to 500Å and plotted the amount of particles as a histogram. For the 3-dimensional analysis, we used the sg_neighbor_plot_local function in STOPGAP ^94^. This function generates a volume where each point represents the position of the nearest neighboring particle, and the density of the point in the volume are integer values corresponding to the number of neighbors found in that position.

#### Scoring of viral morphology

To test the morphology of M VLPs we took montages of cryoEM grid squares (3000x), followed by high magnification (63000x) images. We then scored the VLPs based on their contents. If a particle contained M and another NiV protein (N, F or G), the morphology was assessed and reported as either a sphere, tube, or other. All the samples in this study also contained contaminating vesicles, which were omitted from the analyses. These contaminating vesicles were small, spherical liposomes and did not contain a layer of the M protein along the inner membrane leaflet.

#### Negative Stain TEM

Copper TEM grids were glow discharged at 15mA for 45s. For negative stain, 3uL of VLP containing sample was applied to formvar coated continuous carbon TEM grids. Sample was then stained with 1uL of aqueous 2% Uranyl Acetate for 60s. Excess stain was wicked off and allowed to dry. Images were taken on a Talos F200Ci operating at 120kV equipped with a Ceta CCD detector.

#### CryoET and tomogram reconstruction of transfected cells

5e5 HEK293T cells were seeded on Quantifoil R2/2 Gold 300 mesh grids in 35 mm dishes. Cells were transfected with 3 µg of plasmids encoding NiV structural proteins for 48 hours. Grids were removed from the growth medium and transferred directly to a Vitrobot Mark 3 set to 100% humidity and 25 °C (ThermoFisher Scientific). 3uL of warm PBS was applied to the surface the grid, blotted for 8s and plunge frozen into liquid ethane. Tilt series were acquired on Titan Krios G1 or G4 cryo-electron microscopes (ThermoFisher Scientific). The Titan Krios G1 was equipped with a Gatan K3 direct electron detector operated in counting mode with a BioQuantum energy filter, and data were acquired utilizing a slit width of 20eV. The Krios G4 was equipped with a Falcon 4i direct electron detector and SelectrisX imaging filter, and data were acquired utilizing a slit width of 10ev. Tilt series were acquired using a dose-symmetric tilt scheme (Hagen scheme) from +60° to-60° at 3° increments starting at 0° and reversing direction every 2 tilts ^84^.

Tilt series pre-processing (Import, Motion-Correction and 2-D CTF-estimation) were all carried out using WarpTools to generate tilt-stacks of micrographs at 2Å/pixel ^95^. Tilt stacks were aligned using AreTomo2, and subsequently, transform files and tilt angles were reimported to WarpTools for 3D CTF estimation and tomogram reconstruction. Odd/Even tomograms were also generated to train the cryoCARE neural network for 100 epochs with a learning rate of 0.0004 ^89^.

#### Model Building and Refinement of M-Lattice

An initial model of one monomer of M was generated using AlphaFold2 then the monomer docked into the map using Chimera and modeled in Isolde inside of ChimeraX ^96–98^. Two alpha helices (aa 191-194 and 286-292) were replaced by flexible loops based on the map density observed. The C-terminus of M was rebuilt into density which was consistent with the previously published x-ray crystal structure of the M-Dimer PDB:7SKU ^39^. This monomer was used to build out the full 9-Dimer map using the fit in map function in Chimera ^98^. M-Dimer A and M-Dimer B differed slightly (Fig. 2B and fig. S6). The 9-Dimer model was refined in Phenix using Real Space Refinement with NCS symmetry and secondary structure constraints ^99^.

#### Meoscale modeling and visualization

VLP envelope segmentation was conducted using MemBrain-seg and DragonFly ^100,101^. This process yields three triangulated meshes representing spheres, tubes, and custom shapes. These meshes are subsequently post-processed in Instant Meshes, which automatically retopologized complex meshes into quadrilateral grids ^102^. This step leverages the repetitive quad lattice appearance of M, allowing us to approximate its distribution by remeshing the envelope into quads and positioning F and M at the center of each quad. G is then placed at regions of high curvature (custom shapes) based on segmentation of the tomogram manually. The N assembly is generated through a random walk within the envelope mesh, resulting in approximately 175 N rings, which are further relaxed using NVIDIA Flex. Our modeling approach used the methods developed previously ^103–105^. Due of the lattice distribution, remaining clashes exist between M-M dimers.

Each resulting model is exported as an mmCIF file and accompanied by a manifest file, facilitating integration into *Mol*\* *Explorer* for community sharing ^106^. The final dataset includes both segmented data (envelope volume isosurfaces and individual protein placements with corresponding Star and TBL files) and the integrative model data, providing comprehensive resources for structural and functional analysis for readers to interact with themselves, made publicly available here (Interactive Tour).

#### Quantification of F and G spatial localization on VLP surfaces

Tomograms containing N,M,F and G were denoised with cryoCARE and visualized in IMOD ^86,89^. Snapshots of VLP central slices containing scale bars were taken and imported into Adobe Illustrator. The Adobe Illustrator canvas was set in dimensions of points. A line was drawn on top of the scale bar to determine the conversion between the canvas and the VLP snapshot (1pt = 2.64 nm). Lines were traced along membranes of both “low curvature” and “high curvature” identified by membrane regions with the purple and white boxes in respectively (Fig S12B). We then counted the number of G tetramers and F trimers along that line. Since on average there are more areas of “low curvature” than “high curvature”, we divided the total # of proteins observed per point, and converted to nanometers to normalize the values.

#### Movie S1

Video flying through the Z-axis of an M:N tomogram, highlighting the presence of nucleocapsid assemblies inside the virus-like particle.

#### Movie S2

Video flying through the Z-axis of a tomogram, highlighting the presence of the M:F co-assembly inside a virus-like particle.

#### External URL

An interactive tour meant to supplement this manuscript through the use of molecular models to enhance the communication of our findings. https://molstar.org/me/viewer/?url=https%3A%2F%2Fraw.githubusercontent.com/mesoscope/cellPACK_data/master/models/me_tours/NivTour.molx&type=molx&hide-controls=1j

---

**Figure S1.**
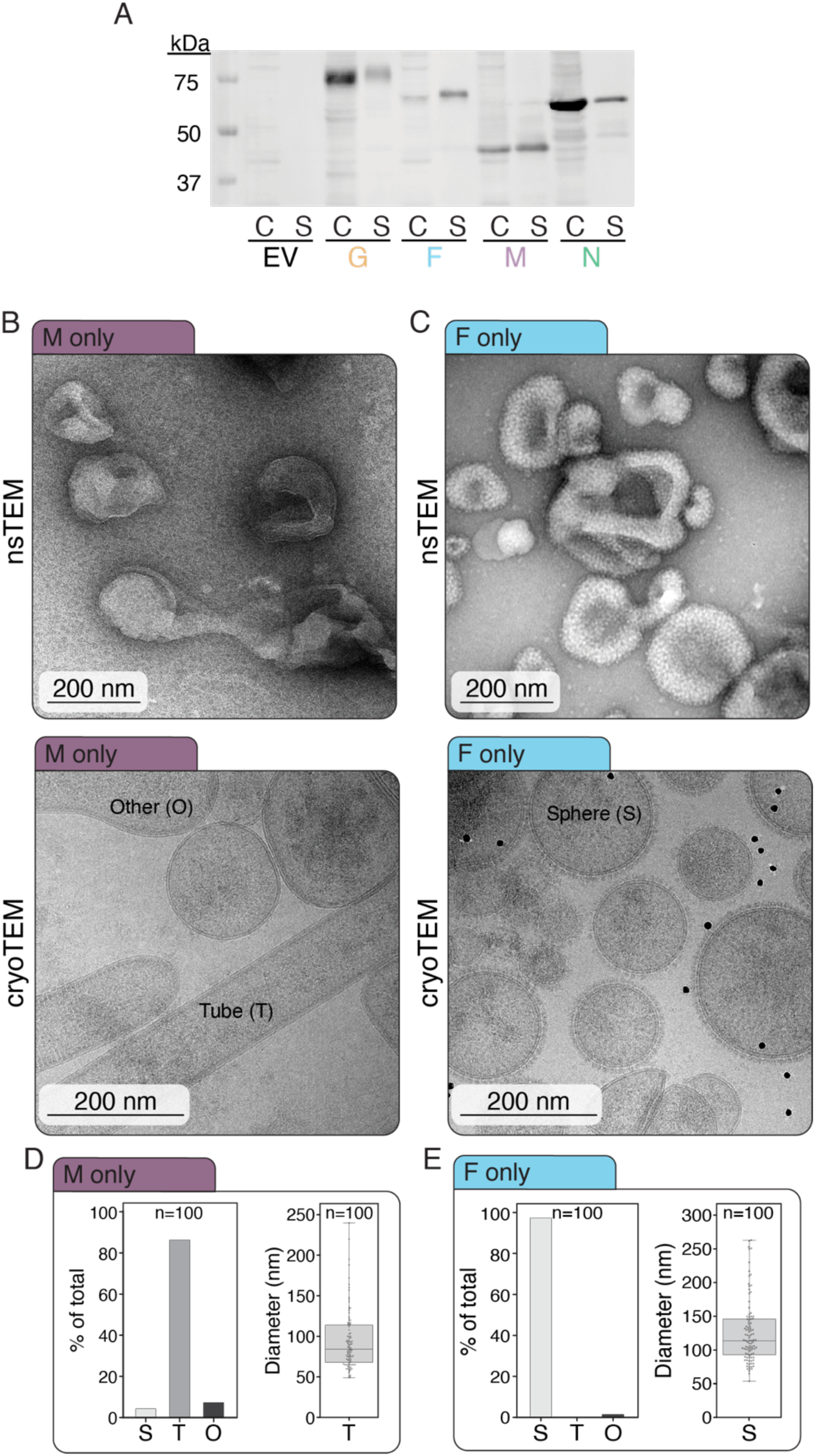
NiV M and F generate VLPs more efficiently than G or N and form distinct morphologies as described by electron microscopy. **(A)** Representative western blot analysis of each individual NiV Protein G (67.1 kDa w/o glycans), F (60.1 kDa w/o glycans), M (39.9 kda), N (58.2 kDa), (C = Cell Lysate), (S = VLP Supernatant) derived from transfected cells 48 hours post transfection with Empty Vector control (EV). **(B)** NiV M-only (left) or **(C)** F-only (right) VLPs imaged with nsTEM (top) or cryoTEM (bottom), annotating the morphologies observed. **(D)** Quantification of morphology (left) and tube diameter (right) of NiV M-only VLPs Mean = 94.7, SD = 36.8. (E) Quantification of morphology (left) and sphere diameter (right) of NiV F-only VLPs Mean = 129, SD = 55.1. S = Sphere, T = Tube, O = Other morphology.

**Figure S2.**
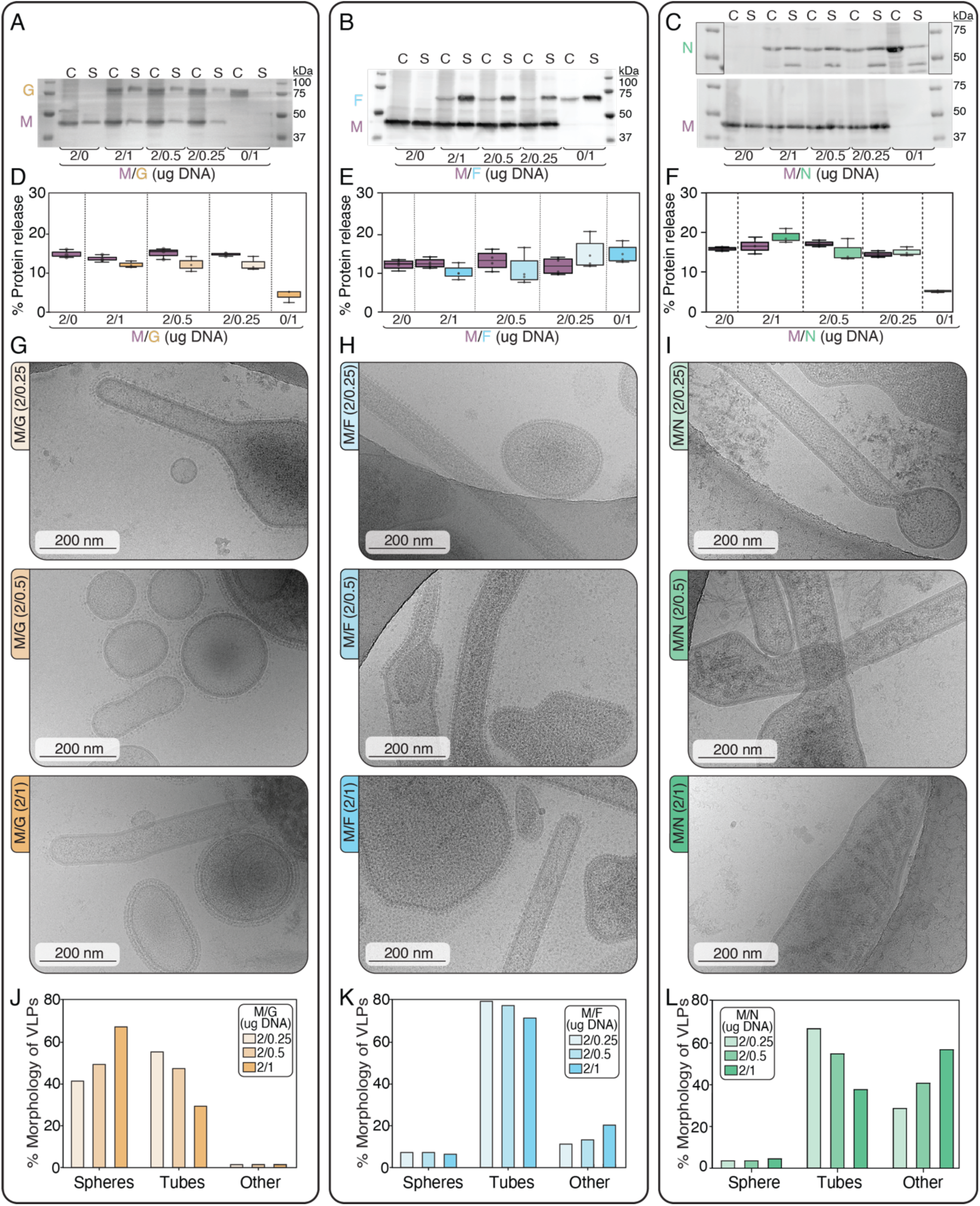
**Analysis of NiV M:G, M:F, and M:N VLPs**. **(A-C)** Representative western blots of NiV M co-transfections with G, F, or N (C=Cell Lysate) (S=VLP Supernatant). **(D-F)** Relative budding efficiency of each individual NiV protein present in co-transfections, calculated by densitometry of total protein present in western blots S/C; n=3. **(G-I)** Representative micrographs of each co-transfection experiment describing the variable morphology present in each combination. **(J-L)** Quantification of particle morphologies present in each co-transfection experiment represented as percentages of at least 50 VLPs per DNA transfection ratio.

**Figure S3.**
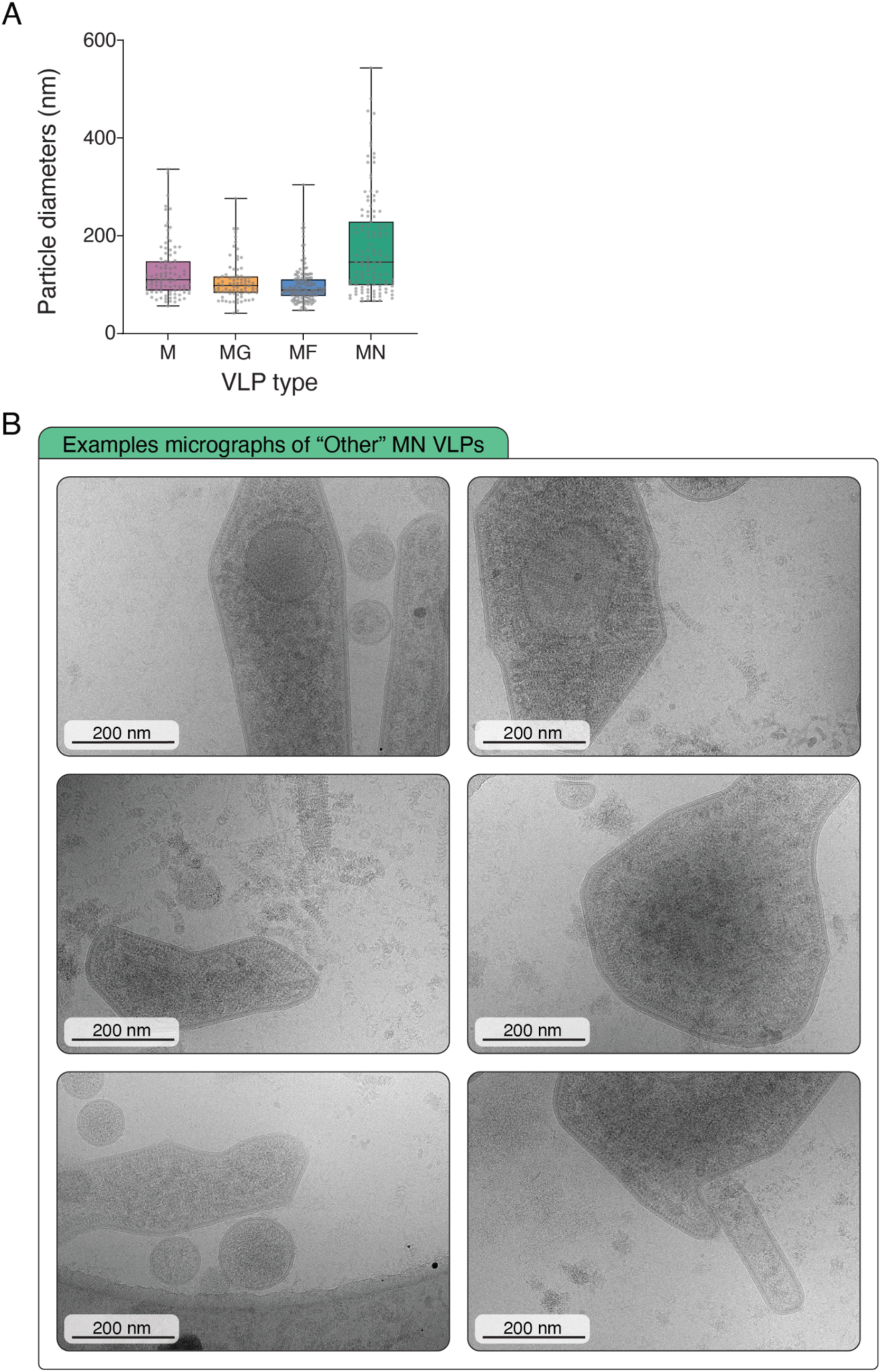
M:N VLP structure and morphology analysis. **(A)** Box plots of the diameters of VLPs from M: G, F or N co-transfections from fig. S2, the largest diameter was measured. **(B)** Representative micrograph gallery of M:N VLPs displaying a pleomorphic phenotype.

**Figure S4.**
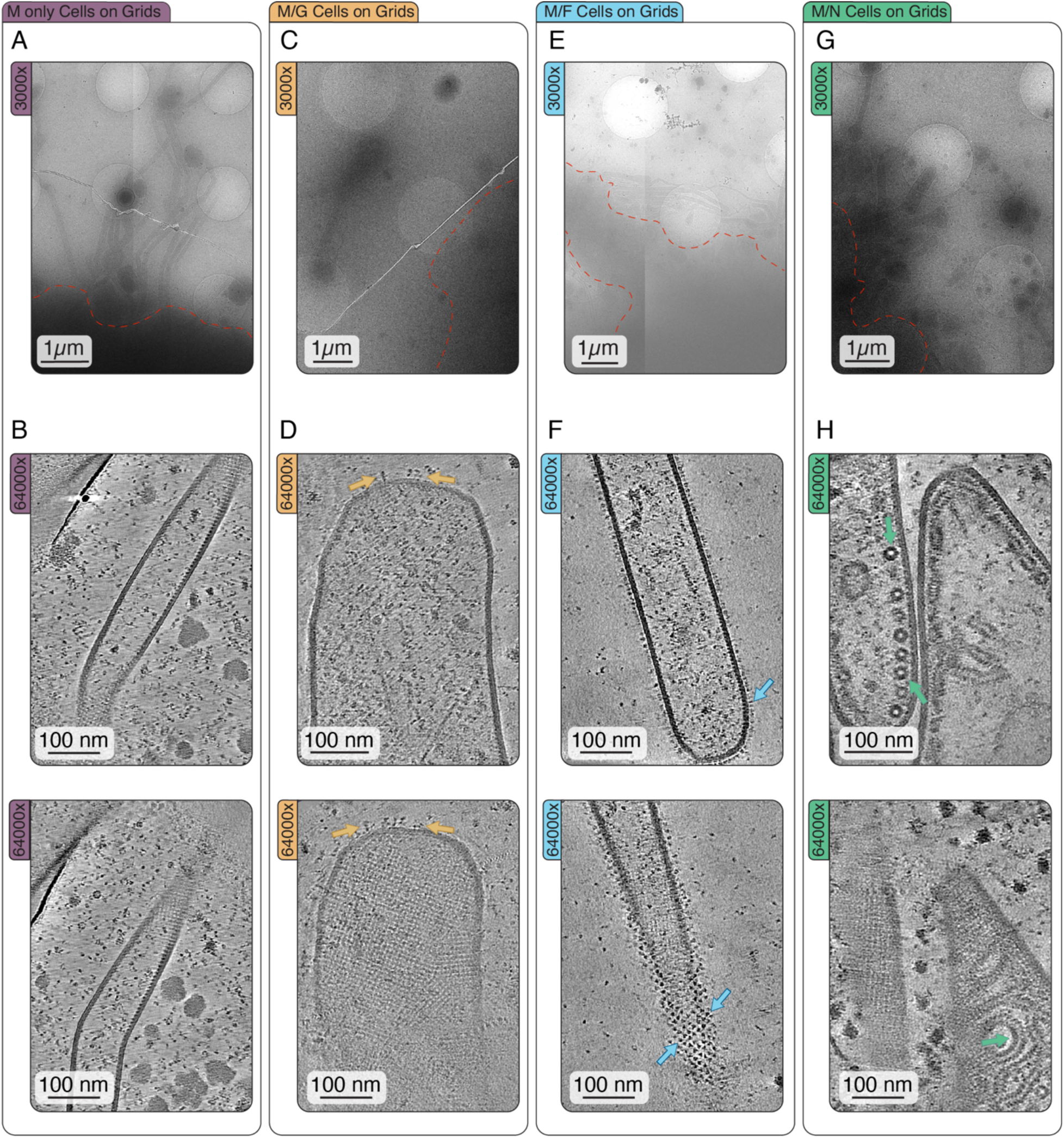
CryoET of VLPs from HEK293T Cells expressing NiV Structural Proteins cultured directly on grids. (A,C,E,G) Low magnification (3000x) of the edges of HEK293T cells, approximate cell edge annotated with a red dashed line, VLPs observed right next to cell edge. **(B,D,F,H)** High magnification tomograms (64000x) of VLPs acquired on or near cell edges, Z slices (10 nm) through the center (top) and M-Lattice (bot) of the same VLP.

**Figure S5.**
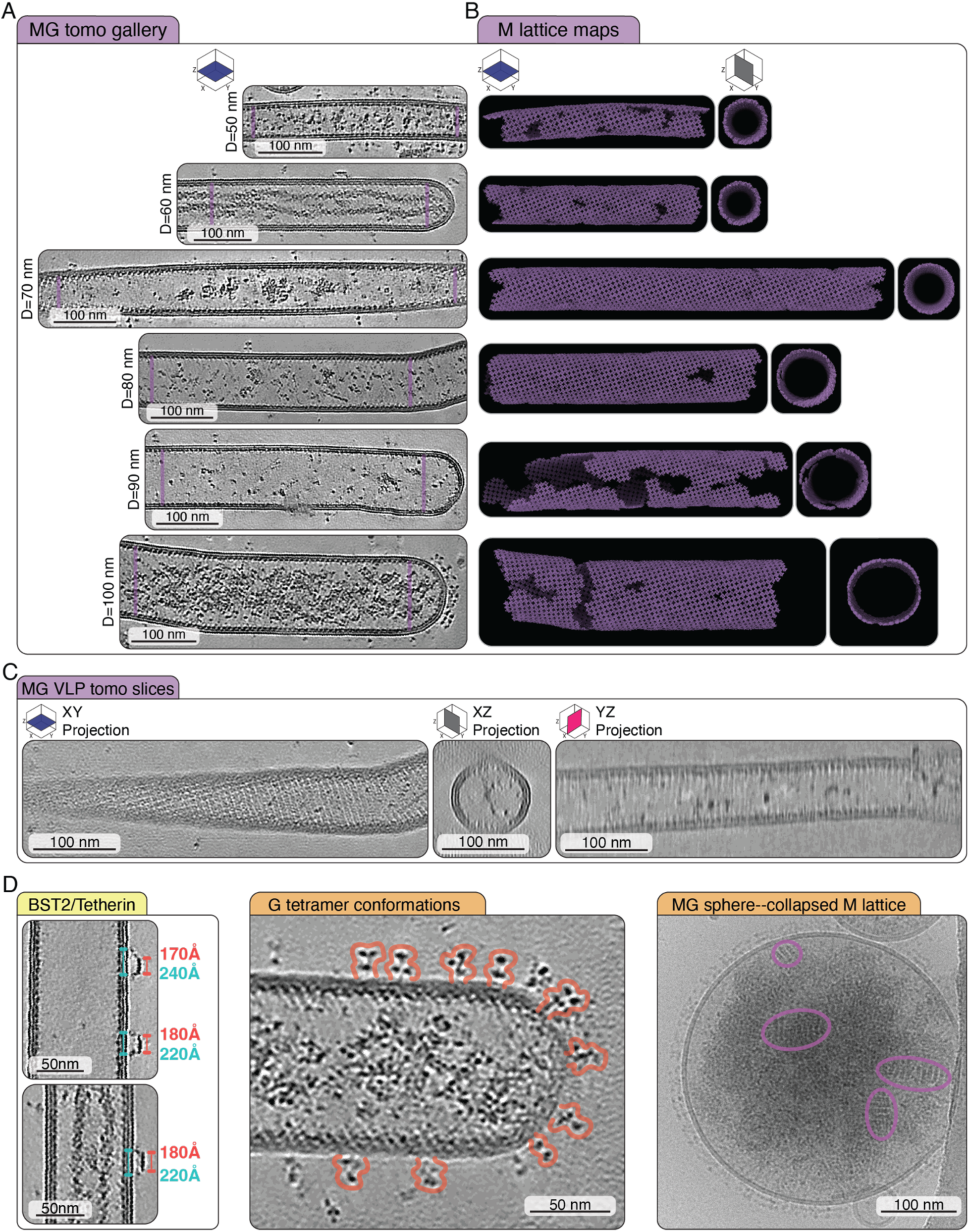
Gallery and analysis of M:G tube VLPs. **(A)** Summed Z-slices from tomograms used in STA of NiV M lattice demonstrating a wide range of tube diameters. **(B)** Lattice mapping information of the aligned subtomogram positions for all the tubes in (A). **(C)** Several projections of a M:G tube used for STA of NiV M. (D) Miscellaneous observations from our M:G VLP data set: Electron microscopy density of a protein with two regions embedded in VLP membrane, measurements consistent with BST2/Tetherin on the sides of tubes (left) ^107–109^, heterogeneous conformations of the G tetramers located on the tips of tubes displaying flexibility of head domains (middle), an example of a spherical VLP containing a high concentration of G on the surface and a collapsed M Lattice inside of the particle (right).

**Figure S6.**
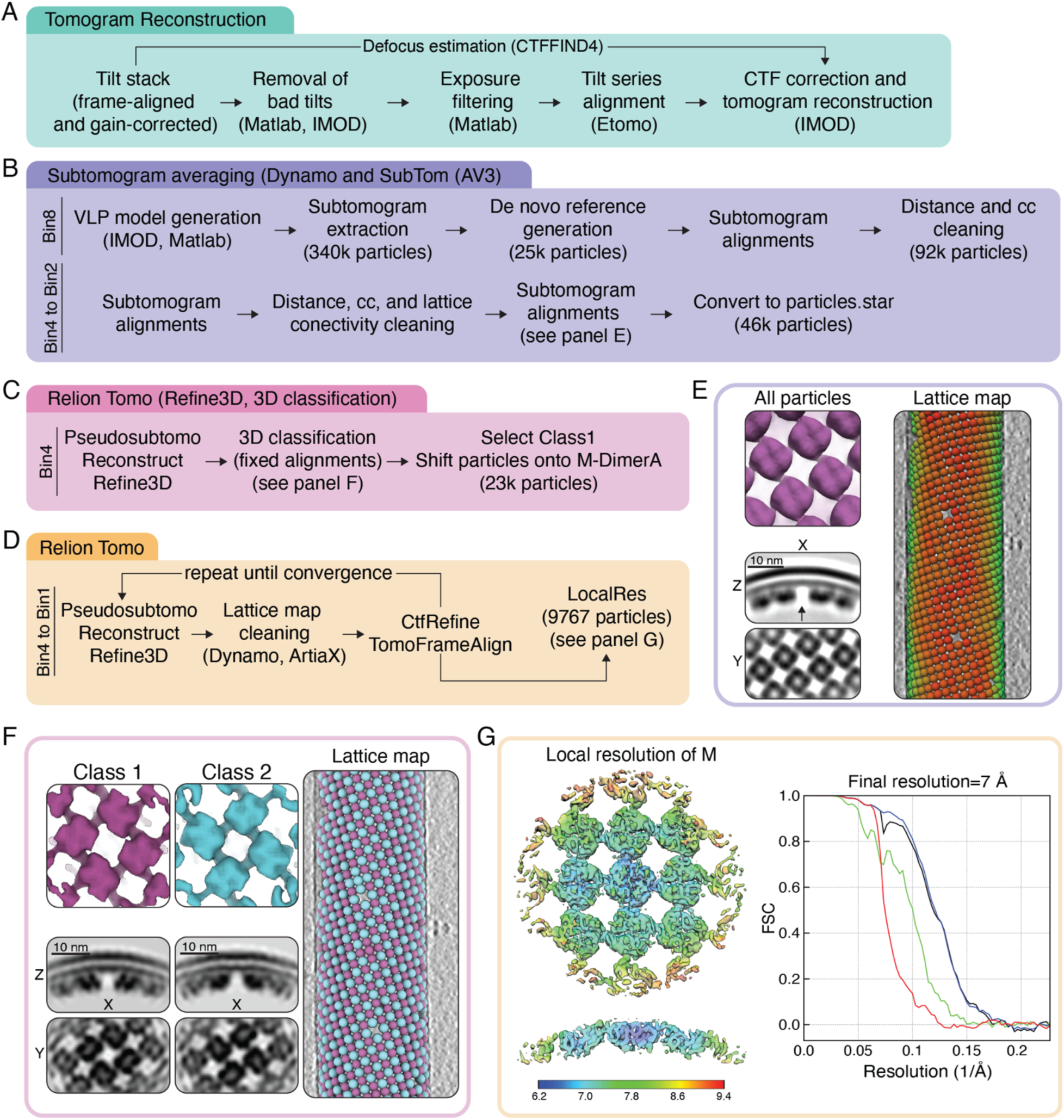
Subtomogram averaging processing pipeline for NiV M Lattice in VLPs. **(A)** Preprocessing. **(B)** Initial low-resolution refinement. **(C)** High-resolution 3-D classification. **(D)** Particle refinement and polishing. **(E)** Bin4 subtomogram average and lattice mapping information of M Lattice before the split of the two classes, coloring based on cc values (red=low cross correlation between subtomogram and template, green = high cross correlation between subtomogram and template). **(F)** Bin 2 3-D classification of NiV M Lattice after the split of the two classes, showing the two tetramers of dimers as isosurface and orthoslice (left) and the reprojections of the two class back onto the lattice map showing the alternation of tetramer classes. **(G)** Final subtomogram average of 9 M Dimers containing both tetramers of dimers (both Class 1 and Class 2) used for modeling, colored by local resolution (left) and FSC curve, estimated resolution cutoff (0.143) (right).

**Figure S7.**
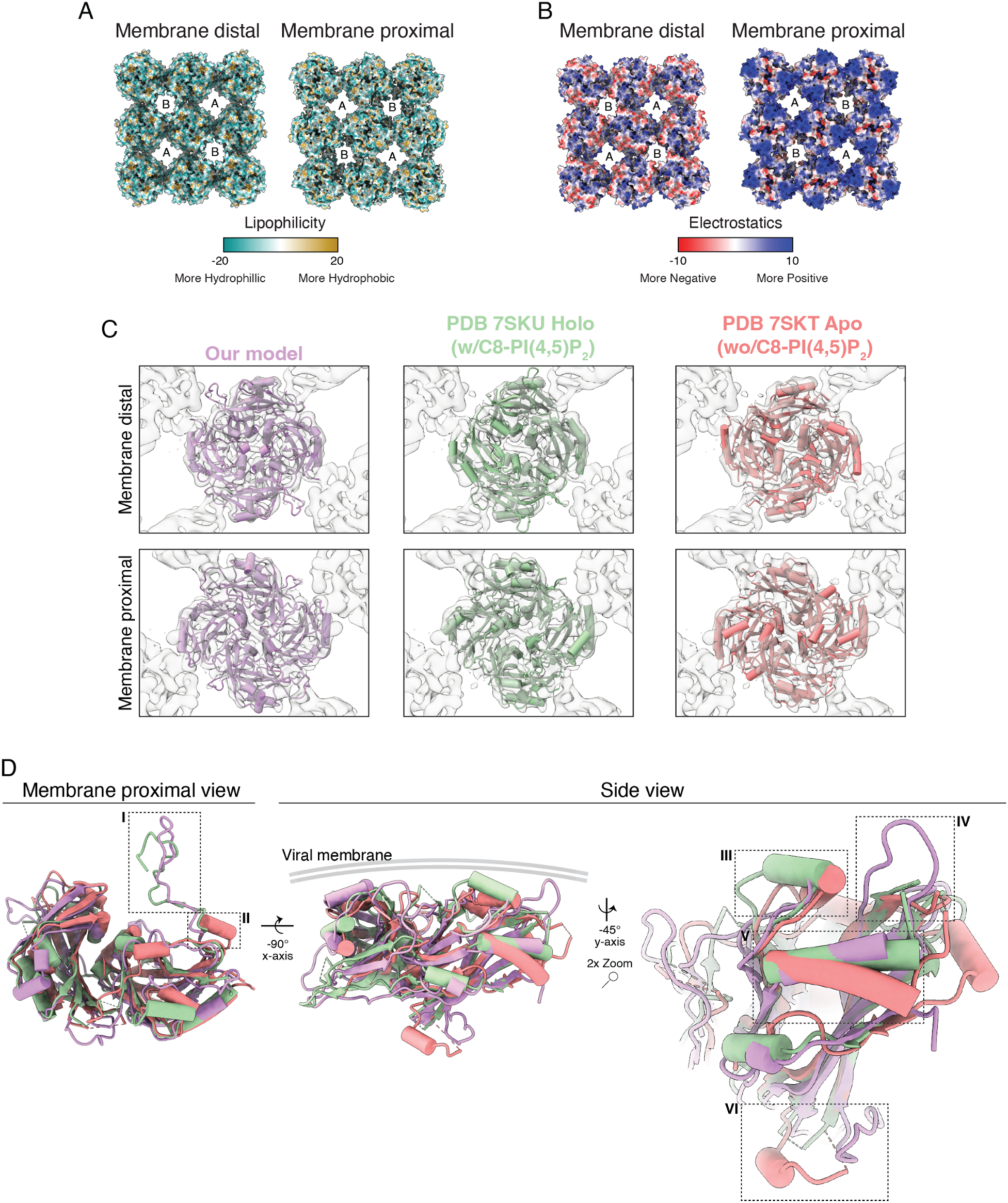
Structural analysis of NiV M. **(A)** Molecular Lipophilicity Potential from the pyMPL implementation in ChimeraX. **(B)** Coulombic electrostatic potential Implementation in ChimeraX. **(C)** Electron microscopy density map of NiV M Lattice from Fig. 2 with atomic models docked, shown as cylinders and stubs PDB: 7SKU (Holo (w/PI(4,5)P_2)_ and PDB 7SKT (Apo (w/o PI(4,5)P_2)_ ^39^. **(D)** Overlay of the atomic models used in (C) displaying several views of the M lattice model. The atomic model for NiV M found in this paper showing resemblances with atomic model PDB: 7SKU (Holo (w/PI(4,5)P_2)_. Similarities and differences between the models are highlighted with boxes (dashed lines) annotated with Roman Numerals I – VI.

**Figure S8.**
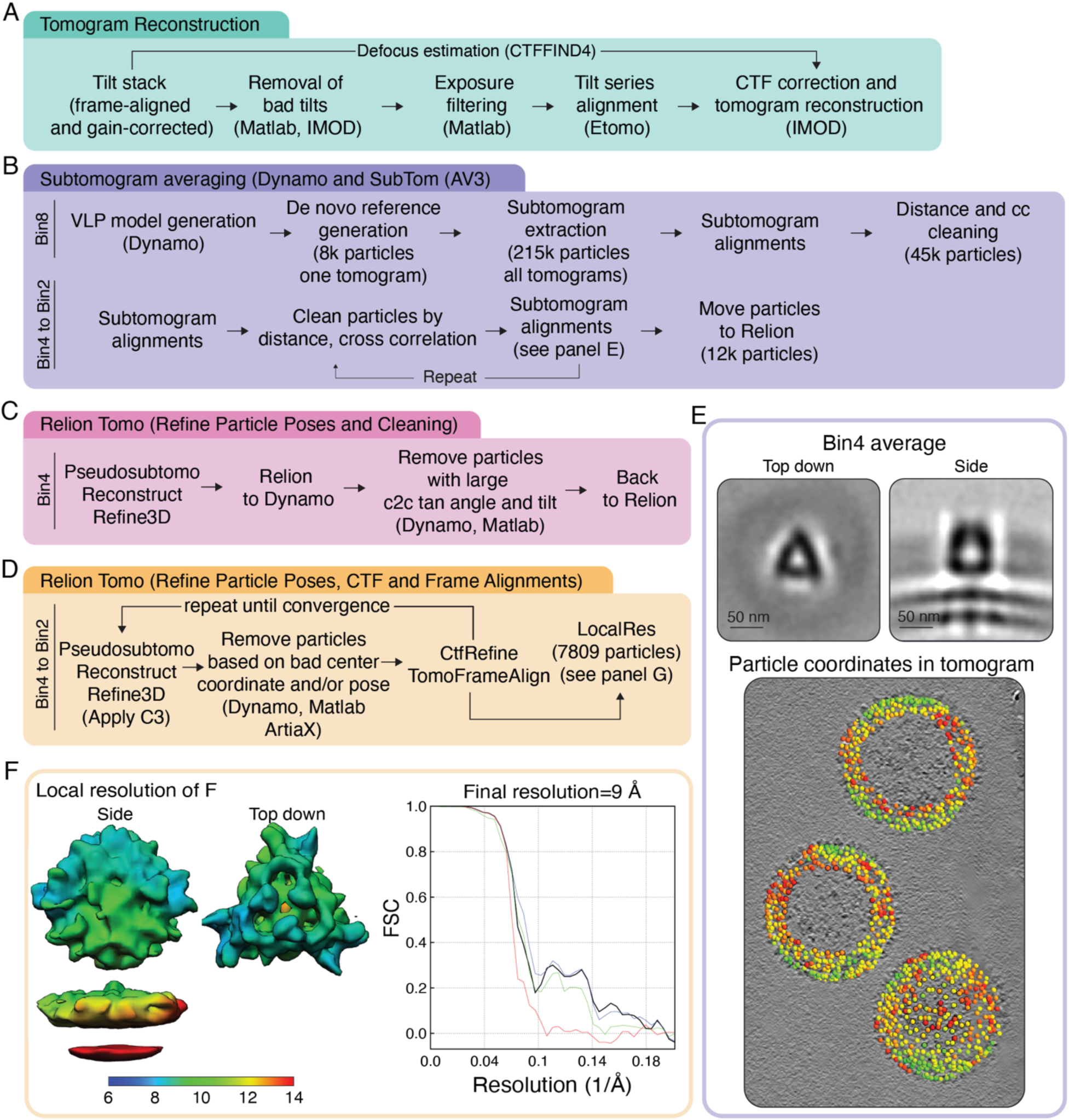
Subtomogram Averaging Processing Pipeline for full-length NiV F on VLPs. **(A)** Preprocessing. **(B)** Initial Low-resolution Refinement. **(C)** High-resolution pose refinement. **(D)** Particle Refinement and Polishing. **(E)** Bin4 subtomogram average and lattice mapping information. **(F)** Local resolution map (w/ gaussian filter for visualization) and FSC with resolution indicated at 0.143 cutoff.

**Figure S9.**
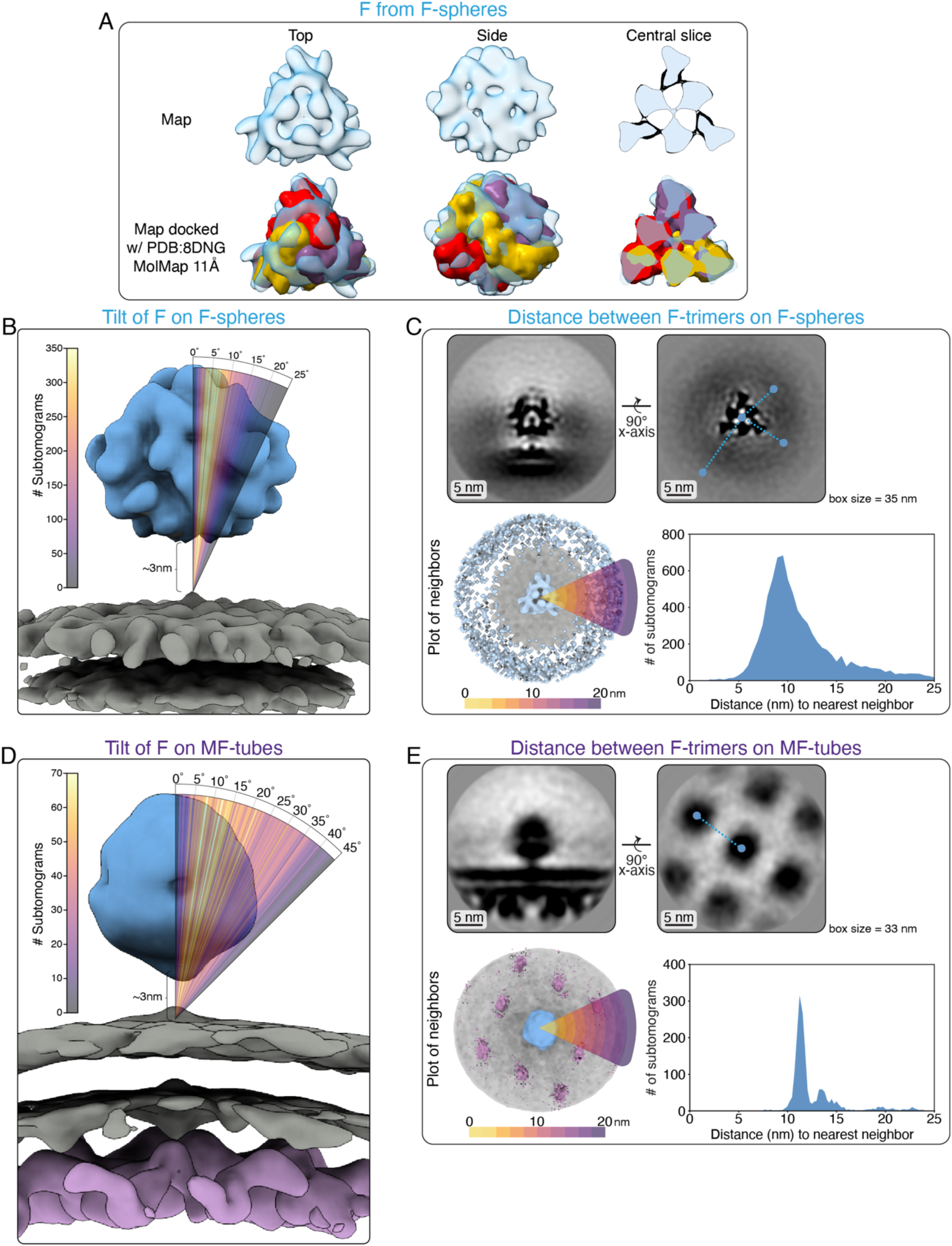
Structural Analysis of full-length NiV F on the surface of F only and M:F VLPs. **(A)** Subtomogram average of full-length NiV F ectodomain (lowpass filter 10Å for depiction) on the surface of VLPs either alone or docked with a MolMap (molmap=map-like density generated from a model at a resolution of 11Å) of PDB 8DNG. **(B)** Tilt of the NiV F ectodomain on F-only VLPs in degrees with respect to the normal angle of the membrane (defined by manually annotating membrane surface). **(C)** Distribution of F on the surface of F-only VLPs. Orthoslice side and top view of F subtomogram average (top). Neighbor plot (bottom left) and distance to nearest neighbor (bottom right) of F on the surface of F-only VLPs. **(D)** Tilt of the NiV F ectodomain on M:F VLPs in degrees with respect to the normal angle of the membrane (defined by M lattice). **(E)** Distribution of F on the surface of M:F VLPs. Orthoslice side and top view of M:F subtomogram average (top). Neighbor plot (bottom left) and distance to nearest neighbor (bottom right) of F on the surface of M:F VLPs.

**Figure S10.**
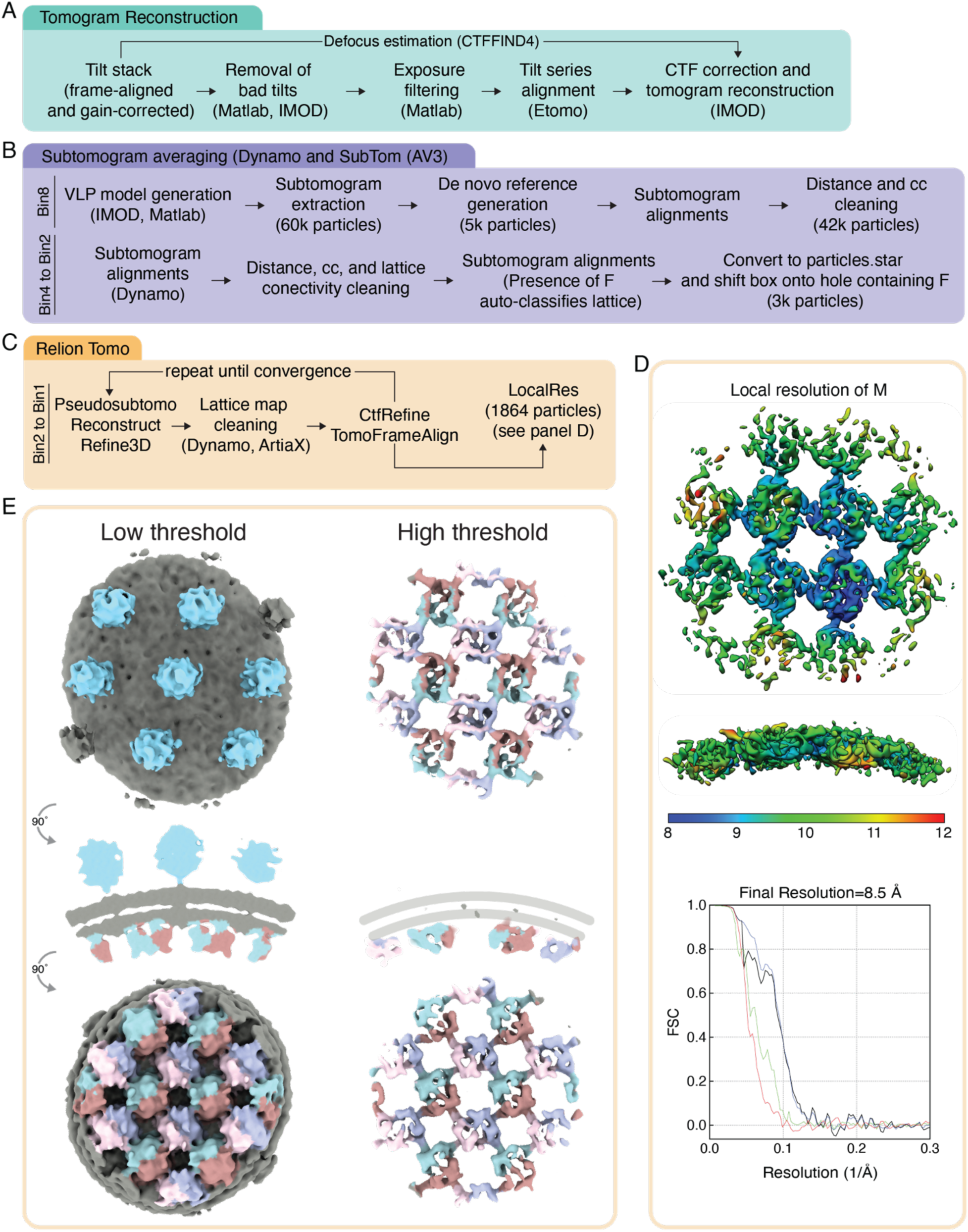
Subtomogram averaging processing pipeline for Nipah M:F complex in VLPs. **(A)** Preprocessing. **(B)** Initial low-resolution refinement. **(C)** High-resolution pose refinement and polishing. **(D)** Local resolution map and FSC resolution cutoff (0.143). **(E)** The same particle positions and tomograms used for (D) re-extracted in a larger box without pose refinement, High and Low threshold, electron microscopy density maps false colored by protein for easier visualization.

**Figure S11.**
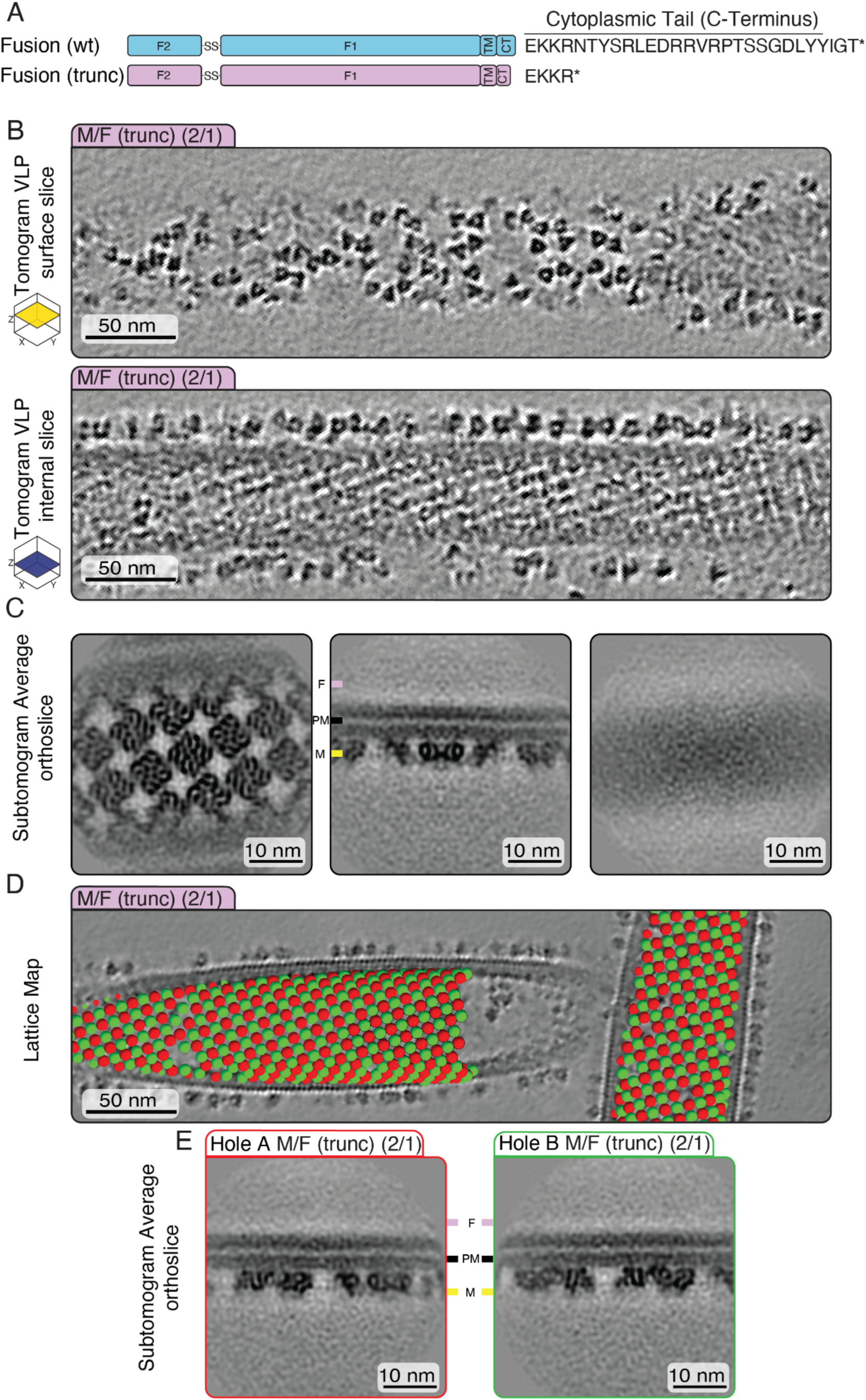
CryoET and subtomogram averaging of NiVM:F(trunc) VLPs. **(A)** Protein schematic of the truncated mutant lacking the majority of the cytoplasmic tail. **(B)** Two Z-slices (10 nm thick) M and F(trunc) VLP tomogram. **(C)** Orthoslice of M:F(trunc) subtomogram averages, focused refinement on M then re extracted into a larger box to visualize F layer. **(D)** Lattice map visualization of the aligned 3-D position and associated class coloring of subtomograms. **(E)** Orthoslices of holes (classes) A and B demonstrating that no F density is present.

**Figure S12.**
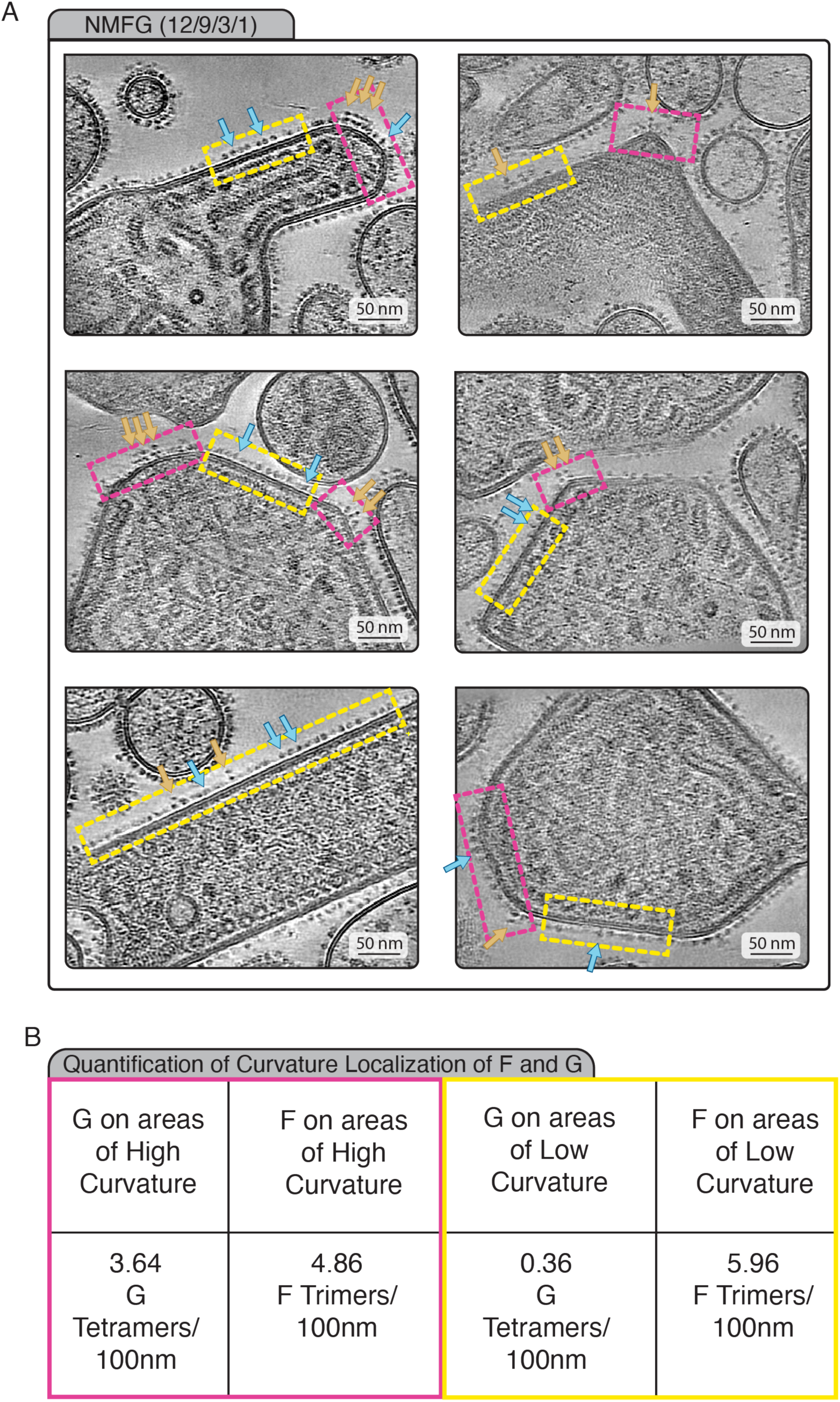
CryoET of and Quantification of NMFG VLPs. **(A)** Snapshots of central z-slices (10 nm thick) taken from tomograms of NMFG VLPs, regions of “low curvature” annotated with yellow dashed line boxes, regions of “high curvature” annotated with pink dashed line boxes. Some examples of F and G are highlighted with blue (F) and orange (G) arrows. **B)** Quantification of the number of G or F oligomers calculated per 100 nm on areas of high vs low curvature.

**Table S1.** CryoEM data collection parameters for Subtomogram Averaging.

| Sample | CryoET of NiV M:G VLPs for Subtomogram Averaging | CryoET of NiV F VLPs for Subtomogram Averaging | CryoET of NiV M:F VLPs for Subtomogram Averaging | CryoET of NiV M:F(trunc)VLPs for Subtomogram Averaging |
| --- | --- | --- | --- | --- |
| Microscope | Talos Arctica | Titan Krios | Talos Arctica | Titan Krios |
| Voltage | 200 keV | 300 keV | 200 keV | 300 keV |
| Energy Filter and Camera | Gatan BioQuantum K3 |  |  |  |
| Energy Filter Slit Width | Yes<br>20eV | Yes<br>20eV | Yes<br>20eV | Yes<br>20eV |
| Å/Pixel | 1.34 | 1.38 | 1.34 | 1.39 |
| Magnification | 63000 | 64000 | 63000 | 64000 |
| Defocus Range | -1.5 to -3.5 | -1.5 to -3.5 | -1.5 to -3.5 | -1.5 to -3.5 |
| Stage Tilt Image Parameters | -60°/60°<br>41 (10 frame) movies | -60°/60°<br>41 (8 frame) movies | -60°/60°<br>41 (10 frame) movies | -60°/60°<br>41 (10 frame) movies |
| Dose Software | 150e<sup>-</sup> Total<br>Serial EM | 150e<sup>-</sup> Total<br>Serial EM | 150e<sup>-</sup> Total<br>Serial EM | 150e<sup>-</sup> Total<br>Serial EM |
| Final Particles | 9767 | 7809 | 1864 | 3768 |
| Final Resolution | M<br>6.8Å (0.143) | F<br>7-13Å | M<br>8.5Å (0.143) | M<br>10Å (0.143) |

**Table S2.**
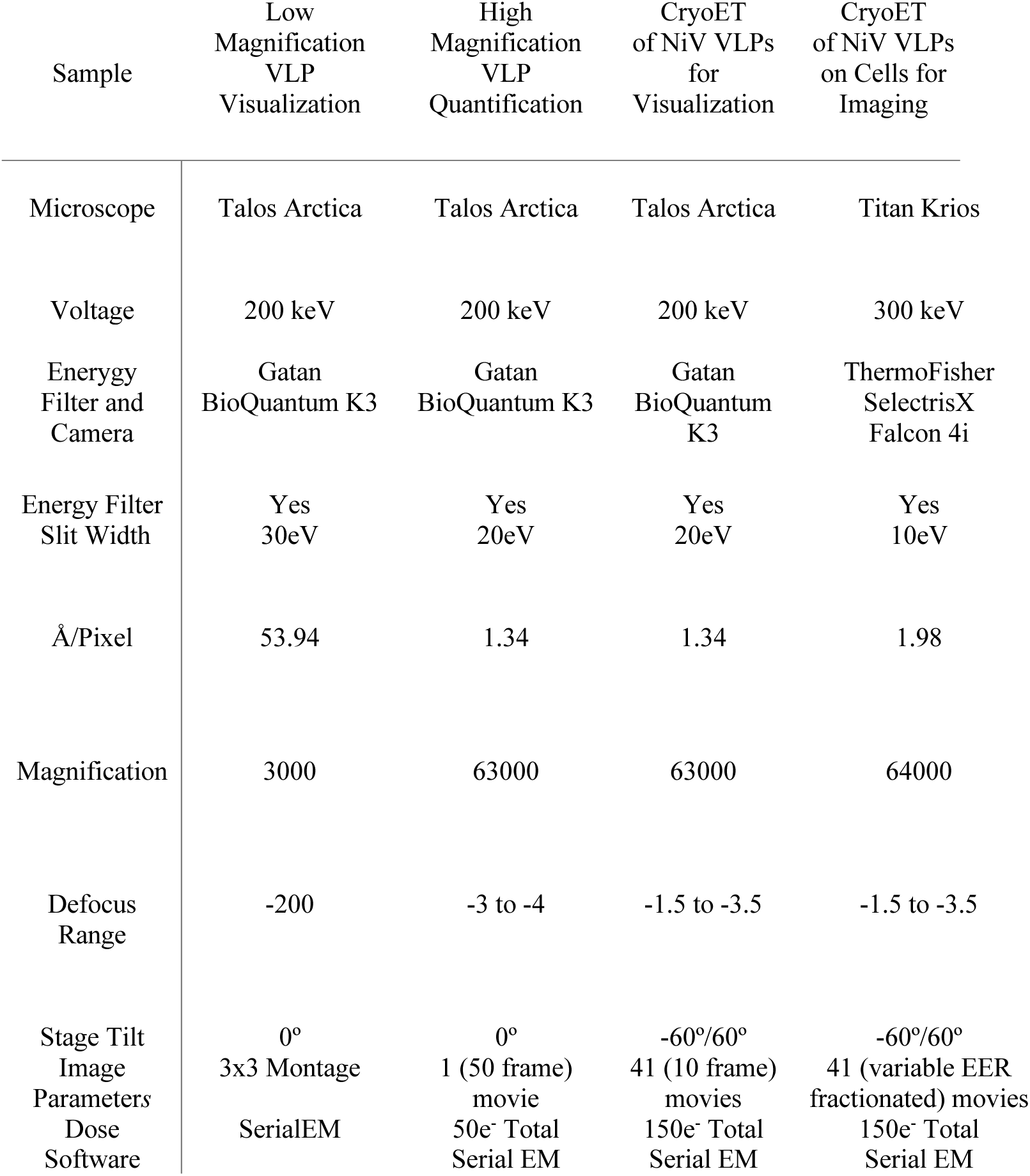
CryoEM data collection parameters for Visualization.

| Sample | Low<br>Magnification<br>VLP<br>Visualization | High<br>Magnification<br>VLP<br>Quantification | CryoET<br>of NiV VLPs<br>for<br>Visualization | CryoET<br>of NiV VLPs<br>on Cells for<br>Imaging |
| --- | --- | --- | --- | --- |
| Microscope | Talos Arctica | Talos Arctica | Talos Arctica | Titan Krios |
| Voltage | 200 keV | 200 keV | 200 keV | 300 keV |
| Energy<br>Filter and<br>Camera | Gatan<br>BioQuantum K3 | Gatan<br>BioQuantum K3 | Gatan<br>BioQuantum<br>K3 | ThermoFisher<br>SelectrisX<br>Falcon 4i |
| Energy Filter<br>Slit Width | Yes<br>30eV | Yes<br>20eV | Yes<br>20eV | Yes<br>10eV |
| Å/Pixel | 53.94 | 1.34 | 1.34 | 1.98 |
| Magnification | 3000 | 63000 | 63000 | 64000 |
| Defocus<br>Range | -200 | -3 to -4 | -1.5 to -3.5 | -1.5 to -3.5 |
| Stage Tilt<br>Image<br>Parameters<br>Dose<br>Software | 0°<br>3x3 Montage<br>SerialEM | 0°<br>1 (50 frame)<br>movie<br>50e<sup>-</sup> Total<br>Serial EM | -60°/60°<br>41 (10 frame)<br>movies<br>150e<sup>-</sup> Total<br>Serial EM | -60°/60°<br>41 (variable EER<br>fractionated) movies<br>150e<sup>-</sup> Total<br>Serial EM |

## Bibliography

1. Chua, K. B. Nipah virus: A recently emergent deadly paramyxovirus. Science (1979). 288, 1432–1435 (2000).

2. Thibault, P. A., Watkinson, R. E., Moreira-Soto, A., Drexler, J. F. & Lee, B. Zoonotic Potential of Emerging Paramyxoviruses: Knowns and Unknowns. Advances in Virus Research vol. 98 (Elsevier Inc., 2017).

3. Chua, K. B. et al. Fatal encephalitis due to Nipah virus among pig-farmers in Malaysia. Lancet 354, 1257–1259 (1999).

4. Ang, B. S. P., Lim, T. C. C. & Wang, L. Nipah Virus Infection. J. Clin. Microbiol. 56, (2018).

5. Luby, S. P., Gurley, E. S. & Hossain, M. J. Transmission of Human Infection with Nipah Virus. Clinical Infectious Diseases 49, 1743–1748 (2009).

6. Arunkumar, G. et al. Outbreak Investigation of Nipah Virus Disease in Kerala, India, 2018. J. Infect. Dis. 219, 1867–1878 (2019).

7. Gurley, E. S. et al. Person-to-person transmission of Nipah virus in a Bangladeshi community. Emerg. Infect. Dis. 13, 1031–1037 (2007).

8. Luby, S. P. et al. Recurrent zoonotic transmission of Nipah virus into humans, Bangladesh, 2001-2007. Emerg. Infect. Dis. 15, 1229–1235 (2009).

9. Russell, C. J., Simões, E. A. F. & Hurwitz, J. L. Vaccines for the Paramyxoviruses and Pneumoviruses: Successes, Candidates, and Hurdles. Viral Immunol. 31, 133–141 (2018).

10. Walpita, P., Barr, J., Sherman, M., Basler, C. F. & Wang, L. Vaccine Potential of Nipah Virus-Like Particles. PLoS One 6, (2011).

11. Coronel, E. C., Murti, K. G., Takimoto, T. & Portner, A. Human Parainfluenza Virus Type 1 Matrix and Nucleoprotein Genes Transiently Expressed in Mammalian Cells Induce the Release of Virus-Like Particles Containing Nucleocapsid-Like Structures. J. Virol. 73, 7035–7038 (1999).

12. Schmitt, A. P., Leser, G. P., Waning, D. L. & Lamb, R. A. Requirements for Budding of Paramyxovirus Simian Virus 5 Virus-Like Particles. J. Virol. 76, 3952–3964 (2002).

13. Pantua, H. D., McGinnes, L. W., Peeples, M. E. & Morrison, T. G. Requirements for the Assembly and Release of Newcastle Disease Virus-Like Particles. J. Virol. 80, 11062–11073 (2006).

14. Li, M. et al. Mumps Virus Matrix, Fusion, and Nucleocapsid Proteins Cooperate for Efficient Production of Virus-Like Particles. J. Virol. 83, 7261–7272 (2009).

15. Liu, Y. C., Grusovin, J. & Adams, T. E. Electrostatic Interactions between Hendra Virus Matrix Proteins Are Required for Efficient Virus-Like-Particle Assembly. J. Virol. 92, (2018).

16. Bracken, M. K. et al. Viral protein requirements for assembly and release of human parainfluenza virus type 3 virus-like particles. Journal of General Virology 97, 1305–1310 (2016).

17. Spehner, D., Drillien, R. & Howley, P. M. The Assembly of the Measles Virus Nucleoprotein into Nucleocapsid-like Particles Is Modulated by the Phosphoprotein. Virology 232, 260–268 (1997).

18. Patch, J. R., Crameri, G., Wang, L.-F., Eaton, B. T. & Broder, C. C. Quantitative analysis of Nipah virus proteins released as virus-like particles reveals central role for the matrix protein. Virol. J. 4, 1 (2007).

19. Johnston, G. P., et al. Nipah Virus-Like Particle Egress Is Modulated by Cytoskeletal and Vesicular Trafficking Pathways: a Validated Particle Proteomics Analysis. mSystems 4, (2019).

20. Johnston, G. P., et al. Nipah Virus-Like Particle Egress Is Modulated by Cytoskeletal and Vesicular Trafficking Pathways: a Validated Particle Proteomics Analysis. mSystems 4, (2019).

21. Battisti, A. J. et al. Structure and assembly of a paramyxovirus matrix protein. Proc. Natl. Acad. Sci. U. S. A. 109, 13996–14000 (2012).

22. El Najjar, F., Schmitt, A. P. & Dutch, R. E. Paramyxovirus glycoprotein incorporation, assembly and budding: a three way dance for infectious particle production. Viruses 6, 3019–3054 (2014).

23. Hyatt, A. D., Zaki, S. R., Goldsmith, C. S., Wise, T. G. & Hengstberger, S. G. Ultrastructure of Hendra virus and Nipah virus within cultured cells and host animals. Microbes Infect. 3, 297–306 (2001).

24. Cox, R. M. & Plemper, R. K. Structure and organization of paramyxovirus particles. Curr. Opin. Virol. 24, 105–114 (2017).

25. Watkinson, R. E. & Lee, B. Nipah virus matrix protein: Expert hacker of cellular machines. FEBS Letters vol. 590 2494–2511 Preprint at 10.1002/1873-3468.12272 (2016).

26. Takimoto, T., Murti, K. G., Bousse, T., Scroggs, R. A. & Portner, A. Role of Matrix and Fusion Proteins in Budding of Sendai Virus. J. Virol. 75, 11384–11391 (2001).

27. Sugahara, F. et al. Paramyxovirus Sendai virus-like particle formation by expression of multiple viral proteins and acceleration of its release by C protein. Virology 325, 1–10 (2004).

28. Pohl, C., Duprex, W. P., Krohne, G., Rima, B. K. & Schneider-Schaulies, S. Measles virus M and F proteins associate with detergent-resistant membrane fractions and promote formation of virus-like particles. Journal of General Virology 88, 1243–1250 (2007).

29. Wang, Z. et al. Architecture and antigenicity of the Nipah virus attachment glycoprotein. Science (1979). 375, 1373–1378 (2022).

30. Dang, H. V. et al. An antibody against the F glycoprotein inhibits Nipah and Hendra virus infections. Nat. Struct. Mol. Biol. 26, 980–987 (2019).

31. Byrne, P. O. et al. Structural basis for antibody recognition of vulnerable epitopes on Nipah virus F protein. Nat. Commun. 14, 1494 (2023).

32. Johnston, G. P. et al. Cytoplasmic Motifs in the Nipah Virus Fusion Protein Modulate Virus Particle Assembly and Egress. J. Virol. 91, (2017).

33. Bloyet, L.-M. The Nucleocapsid of Paramyxoviruses: Structure and Function of an Encapsidated Template. Viruses 13, 2465 (2021).

34. Li, M. et al. Mumps Virus Matrix, Fusion, and Nucleocapsid Proteins Cooperate for Efficient Production of Virus-Like Particles. J. Virol. 83, 7261–7272 (2009).

35. Ray, G., Schmitt, P. T. & Schmitt, A. P. C-Terminal DxD-Containing Sequences within Paramyxovirus Nucleocapsid Proteins Determine Matrix Protein Compatibility and Can Direct Foreign Proteins into Budding Particles. J. Virol. 90, 3650–3660 (2016).

36. Xu, K. et al. Crystal Structure of the Pre-fusion Nipah Virus Fusion Glycoprotein Reveals a Novel Hexamer-of-Trimers Assembly. PLoS Pathog. 11, (2015).

37. Guo, Y. et al. The cryo-EM structure of homotetrameric attachment glycoprotein from langya henipavirus. Nat. Commun. 15, (2024).

38. Ker, D.-S., Jenkins, H. T., Greive, S. J. & Antson, A. A. CryoEM structure of the Nipah virus nucleocapsid assembly. PLoS Pathog. 17, e1009740 (2021).

39. Norris, M. J. et al. Measles and Nipah virus assembly: Specific lipid binding drives matrix polymerization. Sci. Adv. 8, (2022).

40. Ke, Z. et al. Promotion of virus assembly and organization by the measles virus matrix protein. Nat. Commun. 9, (2018).

41. Peukes, J. et al. The native structure of the assembled matrix protein 1 of influenza A virus. Nature | 587, 495 (2020).

42. Sibert, B. S. et al. Assembly of respiratory syncytial virus matrix protein lattice and its coordination with fusion glycoprotein trimers. doi:10.1038/s41467-024-50162-x.

43. Loney, C., Mottet-Osman, G., Roux, L. & Bhella, D. Paramyxovirus Ultrastructure and Genome Packaging: Cryo-Electron Tomography of Sendai Virus. J. Virol. 83, 8191–8197 (2009).

44. Liljeroos, L., Huiskonen, J. T., Ora, A., Susi, P. & Butcher, S. J. Electron cryotomography of measles virus reveals how matrix protein coats the ribonucleocapsid within intact virions. Proc. Natl. Acad. Sci. U. S. A. 108, 18085–18090 (2011).

45. Els, H. J. & Josling, D. Article-Artikel Viruses and Virus-like Particles Identified in Ostrich Gut Contents.

46. Booy, F. P., Ruigrok, R. W. H. & van Bruggen, E. F. J. Electron microscopy of influenza virus. J. Mol. Biol. 184, 667–676 (1985).

47. Byrne, P. O. et al. Structural basis for antibody recognition of vulnerable epitopes on Nipah virus F protein. Nat. Commun. 14, (2023).

48. Avanzato, V. A. et al. A monoclonal antibody targeting the Nipah virus fusion glycoprotein apex imparts protection from disease 2 3. doi:10.1101/2022.09.26.507980.

49. Horikami, S. M., Curran, J., Kolakofsky, D. & Moyer, S. A. Complexes of Sendai virus NP-P and P-L proteins are required for defective interfering particle genome replication in vitro. J. Virol. 66, 4901–4908 (1992).

50. Curran, J., Marq, J. B. & Kolakofsky, D. An N-terminal domain of the Sendai paramyxovirus P protein acts as a chaperone for the NP protein during the nascent chain assembly step of genome replication. J. Virol. 69, 849–855 (1995).

51. Sugita, Y., Noda, T., Sagara, H. & Kawaoka, Y. Ultracentrifugation deforms unfixed influenza A virions. doi:10.1099/vir.0.036715-0.

52. Wong, J. J. W. et al. Monomeric ephrinB2 binding induces allosteric changes in Nipah virus G that precede its full activation. Nat. Commun. 8, 781 (2017).

53. Welch, B. D. et al. Structure of the Parainfluenza Virus 5 (PIV5) Hemagglutinin-Neuraminidase (HN) Ectodomain. PLoS Pathog. 9, (2013).

54. Yuan, P. et al. Structural studies of the parainfluenza virus 5 hemagglutinin-neuraminidase tetramer in complex with its receptor, sialyllactose. Structure 13, 803–815 (2005).

55. Yuan, P. et al. Structure of the Newcastle disease virus hemagglutinin-neuraminidase (HN) ectodomain reveals a four-helix bundle stalk. Proc. Natl. Acad. Sci. U. S. A. 108, 14920–14925 (2011).

56. Norris, M. J. et al. Measles and Nipah virus assembly: Specific lipid binding drives matrix polymerization. Sci. Adv. 8, eabn1440 (2022).

57. Leyrat, C., Renner, M., Harlos, K., Huiskonen, J. T. & Grimes, J. M. Structure and self-assembly of the calcium binding matrix protein of human metapneumovirus. Structure 22, 136–148 (2014).

58. Aguilar, H. C. et al. Polybasic KKR Motif in the Cytoplasmic Tail of Nipah Virus Fusion Protein Modulates Membrane Fusion by Inside-Out Signaling. J. Virol. 81, 4520–4532 (2007).

59. Cifuentes-Muñoz, N. et al. Mutations in the Transmembrane Domain and Cytoplasmic Tail of Hendra Virus Fusion Protein Disrupt Virus-Like-Particle Assembly. J. Virol. 91, (2017).

60. Jenni, S., Horwitz, J. A., Bloyet, L.-M., Whelan, S. P. J. & Harrison, S. C. Visualizing molecular interactions that determine assembly of a bullet-shaped vesicular stomatitis virus particle. doi:10.1038/s41467-022-32223-1.

61. Wan, W. et al. Ebola and Marburg virus matrix layers are locally ordered assemblies of VP40 dimers. Elife 9, (2020).

62. Wang, Q., et al. The nanoscale organization of the Nipah virus fusion protein informs new membrane fusion mechanisms. Preprint at 10.7554/eLife.97017.2 (2024).

63. Byrne, P. O. et al. Prefusion stabilization of the Hendra and Langya virus F proteins. J. Virol. 98, (2024).

64. Avanzato, V. A. et al. A monoclonal antibody targeting the Nipah virus fusion glycoprotein apex imparts protection from disease. Preprint at 10.1101/2022.09.26.507980 (2022).

65. Avanzato, V. A. et al. A monoclonal antibody targeting the Nipah virus fusion glycoprotein apex imparts protection from disease. J. Virol. (2024) doi:10.1128/jvi.00638-24.

66. G.P., J., et al. Cytoplasmic motifs in the nipah virus fusion protein modulate virus particle assembly and egress. J. Virol. 91, 1–13 (2017).

67. Negrete, O. A. et al. EphrinB2 is the entry receptor for Nipah virus, an emergent deadly paramyxovirus. Nature 436, 401–405 (2005).

68. Liu, Q. et al. Nipah Virus Attachment Glycoprotein Stalk C-Terminal Region Links Receptor Binding to Fusion Triggering. J. Virol. 89, 1838–1850 (2015).

69. Porotto, M. et al. The Second Receptor Binding Site of the Globular Head of the Newcastle Disease Virus Hemagglutinin-Neuraminidase Activates the Stalk of Multiple Paramyxovirus Receptor Binding Proteins To Trigger Fusion. J. Virol. 86, 5730–5741 (2012).

70. Liu, Q. et al. Unraveling a Three-Step Spatiotemporal Mechanism of Triggering of Receptor-Induced Nipah Virus Fusion and Cell Entry. PLoS Pathog. 9, (2013).

71. Vahey, M. D. & Fletcher, D. A. Low-Fidelity Assembly of Influenza A Virus Promotes Escape from Host Cells. Cell 176, 281–294.e19 (2019).

72. Li, T. et al. The shape of pleomorphic virions determines resistance to cell-entry pressure. Nat. Microbiol. 6, 617–629 (2021).

73. Lamp, B. et al. Nipah Virus Entry and Egress from Polarized Epithelial Cells. J. Virol. 87, 3143–3154 (2013).

74. Zhang, X. et al. Molecular mechanisms of stress-induced reactivation in mumps virus condensates. Cell 186, 1877–1894.e27 (2023).

75. Segal-Maurer, S. et al. Capsid Inhibition with Lenacapavir in Multidrug-Resistant HIV-1 Infection. New England Journal of Medicine 386, 1793–1803 (2022).

76. Zamora, J. L. R. et al. Third Helical Domain of the Nipah Virus Fusion Glycoprotein Modulates both Early and Late Steps in the Membrane Fusion Cascade. J. Virol. 94, 1–19 (2020).

77. Ithinji, D. G. et al. Multivalent viral particles elicit safe and efficient immunoprotection against Nipah Hendra and Ebola viruses. NPJ Vaccines 7, 166 (2022).

78. Liu, Q., Chen, L., Aguilar, H. C. & Chou, K. C. A stochastic assembly model for Nipah virus revealed by super-resolution microscopy. Nat. Commun. 9, (2018).

79. Johnston, G. P. et al. Cytoplasmic Motifs in the Nipah Virus Fusion Protein Modulate Virus Particle Assembly and Egress. J. Virol. 91, (2017).

80. Mastronarde, D. N. Automated electron microscope tomography using robust prediction of specimen movements. J. Struct. Biol. 152, 36–51 (2005).

81. Kimanius, D., Dong, L., Sharov, G., Nakane, T. & Scheres, S. H. W. New tools for automated cryo-EM single-particle analysis in RELION-4.0. Biochemical Journal 478, 4169–4185 (2021).

82. Rohou, A. & Grigorieff, N. CTFFIND4: Fast and accurate defocus estimation from electron micrographs. J. Struct. Biol. 192, 216–221 (2015).

83. Zheng, S. Q. et al. MotionCor2: Anisotropic correction of beam-induced motion for improved cryo-electron microscopy. Nat. Methods 14, 331–332 (2017).

84. Hagen, W. J. H., Wan, W. & Briggs, J. A. G. Implementation of a cryo-electron tomography tilt-scheme optimized for high resolution subtomogram averaging. J. Struct. Biol. 197, 191–198 (2017).

85. Khavnekar, S., Erdmann, P. S. & Wan, W. TOMOMAN: a software package for large scale cryo-electron tomography data preprocessing, community data sharing, and collaborative computing. bioRxiv (2024) doi:10.1101/2024.05.02.589639.

86. Kremer, J. R., Mastronarde, D. N. & McIntosh, J. R. Computer visualization of three-dimensional image data using IMOD. J. Struct. Biol. 116, 71–76 (1996).

87. Castaño-Díez, D., Kudryashev, M., Arheit, M. & Stahlberg, H. Dynamo: A flexible, user-friendly development tool for subtomogram averaging of cryo-EM data in high-performance computing environments. J. Struct. Biol. 178, 139–151 (2012).

88. Buchholz, T.-O. et al. Chapter 13 - Content-aware image restoration for electron microscopy. in Three-Dimensional Electron Microscopy (eds. Müller-Reichert, T. & Pigino, G. B. T.-M. in C. B.) vol. 152 277–289 (Academic Press, 2019).

89. Buchholz, T. O., Jordan, M., Pigino, G. & Jug, F. Cryo-CARE: Content-aware image restoration for cryo-transmission electron microscopy data. Proceedings - International Symposium on Biomedical Imaging 2019-April, 502–506 (2019).

90. Khavnekar, S. & Wan, W. An approach for coherent periodogram averaging of tilt-series data for improved CTF estimation. bioRxiv (2024) doi:10.1101/2024.10.10.617684.

91. Schur, F. K. M., Hagen, W. J. H., De Marco, A. & Briggs, J. A. G. Determination of protein structure at 8.5Å resolution using cryo-electron tomography and sub-tomogram averaging. J. Struct. Biol. 184, 394–400 (2013).

92. Meng, E. C. et al. UCSF ChimeraX: Tools for structure building and analysis. Protein Science 32, e4792 (2023).

93. Ermel, U. H., Arghittu, S. M. & Frangakis, A. S. ArtiaX: An electron tomography toolbox for the interactive handling of sub-tomograms in UCSF ChimeraX. Protein Sci. 31, e4472 (2022).

94. Wan, W., Khavnekar, S. & Wagner, J. STOPGAP: an open-source package for template matching, subtomogram alignment and classification. Acta Crystallogr. D Struct. Biol. 80, 336–349 (2024).

95. Dimitry Tegunov, A. B. P. N. S. M. W.-M. warpem/warp: v2. 0.0 dev36. Zenodo (2025).

96. Croll, T. I. ISOLDE: A physically realistic environment for model building into low-resolution electron-density maps. Acta Crystallogr. D Struct. Biol. 74, 519–530 (2018).

97. Jumper, J. et al. Highly accurate protein structure prediction with AlphaFold. Nature 596, 583–589 (2021).

98. Pettersen, E. F. et al. UCSF Chimera--a visualization system for exploratory research and analysis. J. Comput. Chem. 25, 1605–1612 (2004).

99. Liebschner, D. et al. Macromolecular structure determination using X-rays, neutrons and electrons: Recent developments in Phenix. Acta Crystallogr. D Struct. Biol. 75, 861–877 (2019).

100. Kass, M., Witkin, A. & Terzopoulos, D. Snakes: Active contour models. Int. J. Comput. Vis. 1, 321–331 (1988).

101. Lorenz, L., et al. MemBrain v2: an end-to-end tool for the analysis of membranes in cryo-electron tomography. BioRxiv (2023).

102. Jakob, W., Tarini, M., Panozzo, D. & Sorkine-Hornung, O. Instant field-aligned meshes. ACM Trans. Graph. 34, (2015).

103. Autin, L., et al. Integrative structural modelling and visualisation of a cellular organelle. QRB Discov. 3, e11 (2022).

104. Klein, T. et al. Instant Construction and Visualization of Crowded Biological Environments. IEEE Trans. Vis. Comput. Graph. 24, 862–872 (2018).

105. Maritan, M. et al. Building Structural Models of a Whole Mycoplasma Cell. J. Mol. Biol. 434, 167351 (2022).

106. Rose, A., Sehnal, D., Goodsell, D. S. & Autin, L. Mesoscale explorer: Visual exploration of large-scale molecular models. Protein Sci. 33, e5177 (2024).

107. Strauss, J. D. et al. Three-Dimensional Structural Characterization of HIV-1 Tethered to Human Cells. J. Virol. 90, 1507–1521 (2016).

108. Swiecki, M. et al. Structural and biophysical analysis of BST-2/tetherin ectodomains reveals an evolutionary conserved design to inhibit virus release. Journal of Biological Chemistry 286, 2987–2997 (2011).

109. Swiecki, M., Omattage, N. S. & Brett, T. J. BST-2/tetherin: Structural biology, viral antagonism, and immunobiology of a potent host antiviral factor. Molecular Immunology vol. 54 132–139 Preprint at 10.1016/j.molimm.2012.11.008 (2013).

